# Monocyte-derived peritoneal macrophages protect C57BL/6 mice against surgery-induced adhesions

**DOI:** 10.1101/2022.07.12.499772

**Authors:** Rinal Sahputra, Krittee Dejyong, Adrian S Woolf, Matthias Mack, Judi Allen, Dominik Ruckerl, Sarah E Herrick

## Abstract

Peritoneal adhesions commonly occur after abdominal or pelvic surgery. These scars join internal organs to each other or to the cavity wall and can present with abdominal or pelvic pain, and bowel obstruction or female infertility. The mechanisms underlying adhesion formation remain unclear and thus, effective treatments are not forthcoming. Peritoneal macrophages accumulate after surgery and previous studies have attributed either pro- or anti-scarring properties to these cells. We propose that there are complex and nuanced responses after surgery with respect to both resident and also monocyte-derived peritoneal macrophage subpopulations. Moreover, we contend that differences in responses of specific macrophage subpopulations in part explain the risk of developing peritoneal scars. We characterised alterations in peritoneal macrophage subpopulations after surgery-induced injury using two strains of mice, BALB/c and C57BL/6, with known differences in macrophage response post-infection. At 14 days post-surgery, BALB/c mice displayed more adhesions compared with C57BL/6 mice. This increase in scarring correlated with a lower influx of monocyte-derived macrophages at day 3 post-surgery. Moreover, BALB/c mice showed distinct macrophage repopulation dynamics after surgery. To confirm a role for monocyte-derived macrophages, we used *Ccr2*-deficient mice as well as antibody-mediated depletion of CCR2 expressing cells during initial stages of adhesion formation. Both *Ccr2*-deficient and CCR2-depleted mice showed a significant increase in adhesion formation associated with the loss of peritoneal monocyte influx. These findings revealed an important protective role for monocyte-derived cells in reducing adhesion formation after surgery.

## Introduction

Peritoneal adhesions commonly occur after abdominal or pelvic surgery and abnormally conjoin internal organs to each other or to the adjacent cavity wall. Adhesions may cause a range of complications, including severe chronic pelvic pain, female infertility, intestinal obstruction, and are a contraindication for future abdominal surgery (1). The peritoneum lines the cavity wall and majority of internal organs, and consists of a surface monolayer of mesothelial cells over a layer of submesothelial connective tissue (2). During surgery, this gliding interface is disrupted and adhesions may develop from fibrin-rich clots that span between injured tissues (3). Several key events, including reduced fibrinolysis (4) and mesothelial-to-mesenchymal transition (MMT) (5, 6), have been shown to be pivotal in adhesion formation. However, cellular and molecular pathways underlying their development are not well defined. Consequently, attempts to prevent adhesions occurring, such as atraumatic surgical technique and use of barrier material, show mixed success and hence are not consistently applied (1). An improved understanding of surgical adhesion formation will lead to better prevention strategies.

Peritoneal macrophages, although known to provide immune surveillance of the abdominal cavity (7), are also involved in the formation of post-surgical adhesions (8-10). Critically, however, there is a lack of consensus as to their exact contribution, with studies attributing both protective as well as pathological roles to macrophages. Experimental expansion of peritoneal macrophages has been shown to have protective effects (9, 11), while somewhat in contradiction, depletion of peritoneal macrophages also reduces adhesion formation (12). The peritoneal cavity contains at least three macrophage subpopulations: a long-lived, self-renewing resident population (F4/80^high^) also known as large peritoneal macrophages, a population derived from monocytes recruited from the blood (F4/80^low^) (13) also known as small peritoneal macrophages, and an intermediate population (F4/80^int^), which consists of monocyte-derived macrophages in the process of differentiating into a cavity resident phenotype (14). Functional differences between these subsets (14-16) and time of analysis may explain the contradictory outcome of studies that have manipulated macrophage numbers. However, which specific subsets are responsible for protective or pathologic properties remains unclear.

Notably, fundamental differences exist between common laboratory mouse strains, BALB/c and C57BL/6, in terms of the response of serous cavity macrophages to infection (17, 18). Furthermore, BALB/c mice have been found to be more susceptible to peritoneal adhesion formation compared with C57BL/6 mice (19). Thus, we hypothesised that the difference in propensity to form adhesions is related to the different profiles of peritoneal cavity macrophages in the different mouse strains. Here, we investigated the dynamics of peritoneal macrophage subsets in BALB/c and C57BL/6 mice after performing surgery to induce adhesions. We found that BALB/c mice were more susceptible to mature peritoneal adhesion formation compared with C57BL/6 mice, but this difference was not apparent until day 14 post-surgery. Surprisingly, adhesion susceptibility in BALB/c mice was associated with a reduced influx of monocyte-derived cells at day 3 after surgery compared with C57BL/6 mice. To confirm a protective role for monocyte-derived macrophages, we analysed surgical adhesion formation in *Ccr2*-deficient mice as well as in monocyte-depleted C57BL/6 mice. These findings revealed an important protective role for monocyte-derived macrophages in reducing mature adhesion formation after surgery.

## Materials and Methods

### Experimental animals

Female BALB/c and C57BL/6 mice, 10 to 12 weeks of age, and approximate weight of 20g, were purchased from a commercial vendor (Envigo, Hillcrest, UK). The *Ccr2-*knock out (KO) colony (originally from The Jackson Laboratory, USA (20)) was backcrossed to a C57BL/6 background for at least 10 generations, bred in-house and shared by J. Grainger (The University of Manchester, UK). Ten-to twelve-week-old male and female KO mice and their wild-type (WT) littermates were used. All animals were maintained in individually ventilated cages under specific-pathogen-free conditions in the Biological Services Facilities (BSF) of the University of Manchester and provided with food and water *ad libitum*. Animal experiments were performed according to the UK Animals (Scientific Procedures) Act (1986) under a Project License (P1208AD89) granted by the UK Home Office and approved by the University of Manchester Ethical Review Committee.

### Surgical procedure

Abdominal surgery was performed following a previously described protocol with slight modification (21, 22). Briefly, mice were anesthetized with a mixture of inhaled isoflurane and oxygen, received s.c. injection of buprenorphine analgesia (50-100µl/kg), were shaved and the surgery site cleaned with sterile wipes. All surgical procedures were performed in a dedicated surgical theatre. A midline incision was made through the abdominal wall and peritoneum. Circular wounding, approximately 7 mm^2^ of the peritoneal cavity wall (approximately 1 cm caudal to the xiphoid cartilage and 1 cm from the midline incision) was created using a trauma instrument developed by Dr. M. Eastwood (Department of Biomedical Sciences, University of Westminster, London, UK). The instrument comprised a clamping device that allowed the trauma to be size- and site-specific, and an abrading rod with collar that restricted the depth of insertion and hence pressure applied (supplementary Figure 1). The inserted rod was rotated three times to produce a consistent lesion on the peritoneal cavity wall. Thereafter, approximately 1 cm length of caecum serosa was abraded with a No 15 surgical blade thirty times and then closely opposed to the wounded peritoneal wall by placing two simple interrupted sutures (8-0 Polyamide, non-absorbable, ETHILON W8170, Ethicon, New Jersey, US) horizontally 3-5 mm either side from the peritoneal wound. Linea alba and skin were closed separately using 6-0 non-absorbable monofilament polyamide sutures (ETHICON W1600T, Ethicon, New Jersey, US). A graphical illustration of the experimental approach and a photo of the trauma instrument are provided in supplementary Figure 1. The mice were necropsied by CO_2_ asphyxiation in a rising concentration at day 3, day 7, and day 14 post-surgery.

### Antibody depletion of monocytes

To assess the importance of monocyte-derived peritoneal macrophages during surgery-induced peritoneal adhesion formation, female C57BL/6 mice were treated with 20 μg of α-CCR2 mAb (clone: MC21, RRID:AB_2314128; (23) or isotype control antibody (Rat IgG2b, Biolegend) in 200 μl of PBS via daily intra-peritoneal (i.p.) injection at day -1, 0, and day 1 post-surgery.

### Peritoneal lavage and adhesion score measurement

Collection of peritoneal lavage fluid and peritoneal exudate cells (PECs) was performed by flushing the peritoneal cavity with 3 ml ice cold RPMI 1640 media (Sigma-Aldrich, Dorset, UK) containing 1 % Penicillin/Streptomycin (Sigma-Aldrich, Dorset, UK) 3 times. The first 3 ml of the wash was centrifuged, and the supernatant used for the detection of soluble mediators using ELISA. The cell pellet left was mixed with the remaining washes and PECs collected by centrifugation for flow cytometry analysis. To remove erythrocytes, cells were incubated with red blood cell lysis buffer (Sigma-Aldrich) following the manufacturer’s instructions. Cell number was determined using a Countess II FL automated cell counter (Invitrogen). Adhesion profile was assessed by two independent observers to determine the total number of adhesions within the peritoneal cavity after surgery and also an arbitrary adhesion score. The adhesion score was measured based on the type of adhesion (supplementary Table 1). The more distal the adhesion from the injury site, the higher the adhesion score. The observers were blinded to the treatment groups during the adhesion score assessment but not to the strain of mouse. Adhesion tissue was collected and fixed in 4% buffered paraformaldehyde overnight and then stored in 70% EtOH before being processed and embedded in paraffin wax for histology analysis.

### Flow cytometry

An aliquot (1×10^6^) of total PECs were stained using a viability assay (Zombie UV, Biolegend) and blocked with 5 μg/mL anti-CD16/32 (clone 93; Biolegend) and heat inactivated normal mouse serum (1:10, Sigma-Aldrich) in flow cytometry buffer (0.5% BSA and 2 mM EDTA in Dulbecco’s PBS) before surface staining on ice with antibodies listed in supplementary Table 2 and 3. For intracellular staining, samples were fixed and permeabilised using eBioscience Foxp3 Staining Buffer kit (Fisher Scientific) following the manufacturer’s instructions before adding antibodies in 1x eBioscience™ Permeabilization Buffer (Thermo Fisher Scientific). All antibodies were purchased from Biolegend unless stated otherwise. PECs were analysed on a BD FACSymphony machine using BD FACSDiva software (BD Biosciences) and post-acquisition analysis performed using FlowJo v10 software (BD Biosciences). The number of cells per cavity for each population (e.g. F4/80^high^ resident macrophages) was calculated by determining the percentage of single live cells for each population using FlowJo multiplied by the total number of PEC divided by 100. The gating strategy for flow cytometry analysis is shown in supplementary Figure 2 and 3.

### Histological analysis

Serial sections (5-μm-thick) of peritoneal adhesion tissue were collected, de-waxed and rehydrated using serial dilutions of alcohol prior to Masson’s trichrome staining following manufacturer’s instruction (Sigma-Aldrich). Images were acquired using a [20×/0.80 Plan Apo] objective using the Pannoramic 250 Flash II slide scanner (3D Histech Ltd., Hungary). Collagen profile was analysed by quantifying the selected site-specific adhesion area (supplementary Figure 4) in µm^2^ using RGB thresholding by QuantCentre and HistoQuant plugin on SlideViewer software (version 2.5).

### Detection of soluble mediators

Interleukin (IL)-6, IL-10, IL-12, IL-13, TNF-α, IFN-γ, Relmα and Ym1 in peritoneal lavage fluid were detected by enzyme linked immunosorbant assay (ELISA). Ninety-six-well immunoGrade plates (BrandTech Scientific, Inc) were coated with primary antibodies diluted in appropriate buffer at 4°C overnight. After washing and non-specific binding blocking, the wash supernatant and standard were added, and the samples were incubated at 4°C overnight. Plates were washed and biotinylated detection antibody was added before the plates were incubated with Streptavidin peroxidase (1:1,000, Sigma) for 60 min. Plates were washed and then the TMB substrate solution (Biolegend) was added. After adding stop solution, plates were read at 450 nm on a VersaMax Microplate reader (Molecular Devices). List of coating and detection antibodies is shown in supplementary Table 4.

### Statistical analysis

Statistical analysis was performed using Prism 9 (Graph-Pad software Inc., La Jolla, CA). Data were tested for normal distribution using a Shapiro-Wilk test prior to analysis of significant differences between groups. Data deemed to follow Gaussian distribution were subsequently analysed using t-test for analysis of two groups, or ANOVA followed by Tukey or Sidak posthoc test for experiments with more than 2 groups. Data with non-Gaussian distribution were analysed using Mann Whitney test (2 groups) or Kruskal-Wallis ANOVA followed by Dunn’s multiple comparison test for experiments with more than 2 groups. Differences with a p-value of less than 0.05 were considered statistically significant.

## Results

### Early monocyte influx in C57BL/6 mice is associated with fewer peritoneal adhesions

To determine whether BALB/c and C57BL/6 mice showed differences in susceptibility to peritoneal adhesions and if this related to variations in macrophage subset responses, mice of each strain were subjected to surgical injury. Three, 7 and 14 days after surgery, animals were analysed for adhesion formation between the injured sites on the caecum and the peritoneal cavity wall (site-specific) as well as to other organs or the midline incision (non-site-specific) (see scoring system, supplementary Table 1). Similar to previously published data (19), we found that BALB/c mice were more likely to develop an increased number of peritoneal adhesions as well as involving other organs compared with C57BL/6 mice. However, in our model this enhanced susceptibility did not become apparent until day 14 after surgery (Figure 1A, B). Collagen profile in site-specific adhesion tissue, as assessed by Masson’s trichrome staining, was similar between the two strains at day 3 and 7 post-surgery (supplementary Figure 5). However, BALB/c mice showed significantly higher collagen deposition in the site-specific adhesion tissue at day 14 (Figure 1C, D).

**Figure 1.**
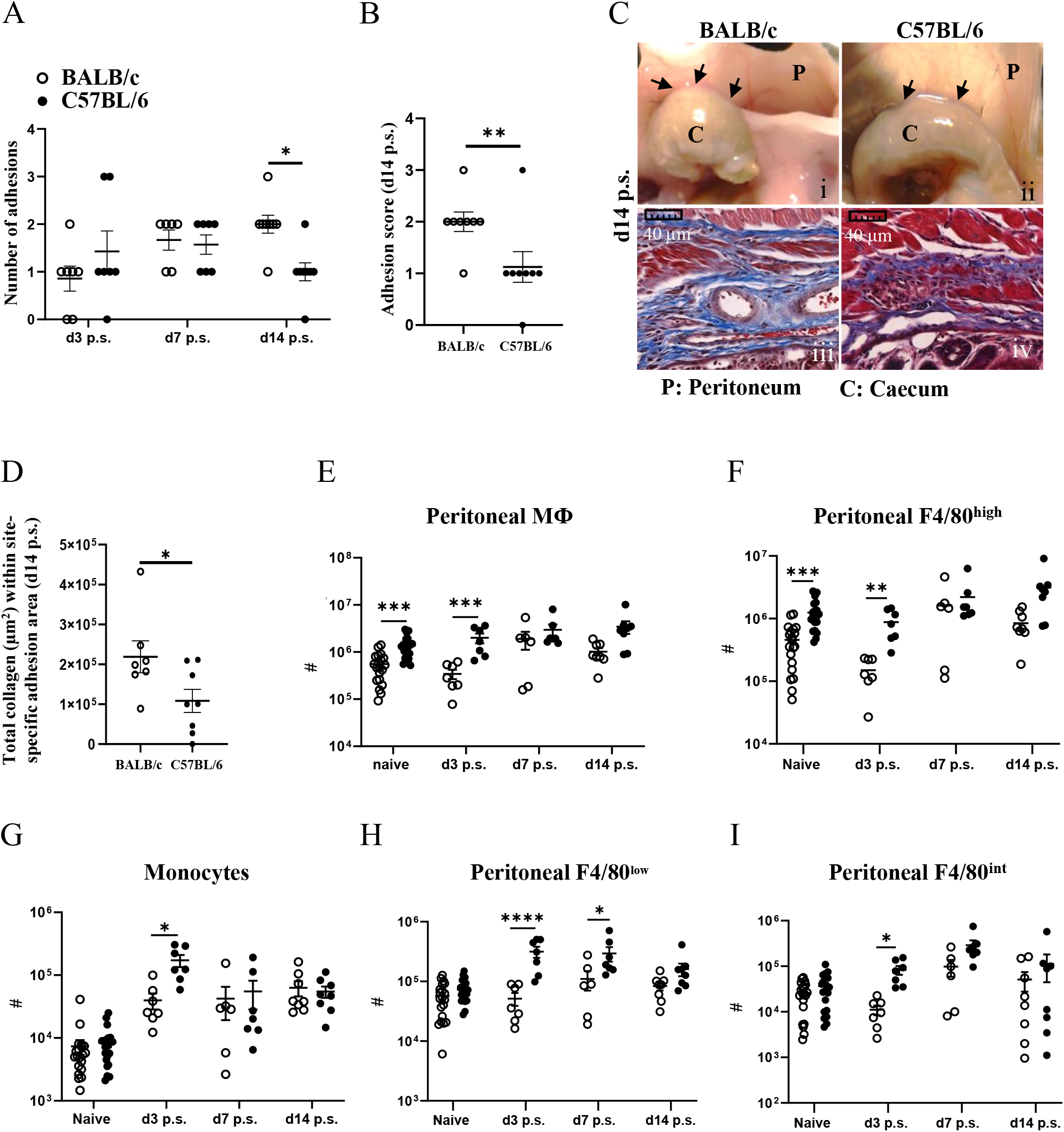
BALB/c mice showed more peritoneal adhesions than C57BL/6 mice at day 14 post-surgery (p.s.), and fewer monocytes and macrophages at day 3. Number of adhesions after surgery at different time points(A) and adhesion score at day 14 post-surgery (B). Representative image and corresponding histological appearance (Masson’s trichrome staining) of adhesions between mouse strains at day 14 after surgery(C). Collagen analysis in adhesion histological sections at day 14post-surgery (D). Total cell number of total peritoneal cavity macrophages (E), peritoneal resident macrophages (F), monocytes (G), peritoneal F4/80^low^ macrophages (H) and peritoneal F4/80^int^ intermediate macrophages (I). Data shows mean±SEM, pooled from 6 separate experiments for naïve or 2 separate experiments for day 3, 7, and 14 post-surgery., n= 6-20 mice/group; *P<0.05, **P<0.01, ***P<0.001, ****P<0.0001. (A) ANOVA with Sidak’s multiple comparison test; (Band C) Mann Whitney test and (E-I) ANOVA with Tukey’s multiple comparison test. Scale bar: 40μm

To assess whether this difference in adhesion formation was associated with variation in macrophage profiles, we analysed PECs by flow cytometry using panels (supplementary Table 2 and 3) that allow us to characterise different subsets of peritoneal macrophages. Although there was no significant difference in the total number of PECs between BALB/c and C57BL/6 prior to and after surgery (supplementary Figure 6), BALB/c mice showed significantly lower number of total peritoneal macrophages (CD11b+Lin-Ly6C-) under steady state conditions as well as on day 3 post-surgery (Figure 1E). This difference was largely due to reduced numbers of cavity resident macrophages (F4/80^high^), which however reached similar numbers in both strains from day 7 (Figure 1F). Moreover, although both strains of mice showed recruitment of monocytes (CD11b+Lin-MHCII-F4/80-Ly6C^high^) from day 3 post-surgery with the number staying elevated up to day 14 post-surgery, the early influx of monocytes (day 3 post-surgery) was significantly more pronounced in C57BL/6 compared with BALB/c mice (Figure 1G). Infiltrating monocytes have been shown to differentiate into mature macrophages (F4/80^low^) within the cavity (24). Moreover, recruited monocytes can also give rise to resident macrophages (F4/80^high^) via an intermediate population (F4/80^int^) (14). In line with the enhanced early influx of monocytes, C57BL/6 mice also showed an enhanced total cell number of F4/80^low^ (Figure 1H) and F4/80^int^ macrophages (Figure 1I) at day 3 post-surgery. In summary, all peritoneal macrophage subsets (F4/80^low^, F4/80^int^, F4/80^high^) were significantly less abundant in BALB/c mice compared with C57BL/6, particularly at day 3 post-surgery and this was associated with enhanced adhesions with increased collagen deposition in BALB/c mice at day 14 after surgery.

### C57BL/6 mice show altered macrophage phenotypes in line with enhanced monocyte-to-F4/80^high^ macrophage conversion

Changes in macrophage dynamics, in particular the loss of F4/80^high^ cavity resident macrophages after surgery, has previously been reported with C57BL/6 mice (25). However, specific differences between BALB/c and C57BL/6 mice have not been investigated; hence we performed a more in-depth characterisation of the dynamic changes in peritoneal macrophage subsets following surgery in the two strains. CD102, and the transcription factor Gata6 are known as defining markers of F4/80^high^ cavity resident macrophages, upregulated at different stages of the differentiation process (26). Both markers were expressed at similar levels by F4/80^high^ macrophages present in both BALB/c and C57BL/6 mice (Figure 2A, B), indicating that although surgery induced considerable changes in the number of these cells with reduced number in BALB/c at day 3 post-surgery (Figure 1F), they retained their peritoneal cavity resident identity. However, at day 3 after surgery F4/80^high^ macrophages of both strains showed a dramatic loss in the proportion of cells expressing Tim4, another defining marker of F4/80^high^ cavity resident macrophages (Figure 2C). Of note, F4/80^high^ macrophages in BALB/c mice regained Tim4 expression over the course of the study with expression at day 14 post-surgery similar to naïve mice (Figure 2C). In contrast, this recovery of Tim4+ F4/80^high^ cells did not occur in C57BL/6 mice (Figure 2C). Expression of Tim4 by F4/80^high^ macrophages has been implicated as a marker of an embryonic origin i.e. established resident peritoneal population, whereas monocyte-derived resident macrophages largely fail to express Tim4 (14). Thus, the persistent loss of Tim4 expression in F4/80^high^ cavity macrophages of C57BL/6 mice might indicate an enhanced integration of monocyte-derived cells into the resident macrophage pool.

**Figure 2.**
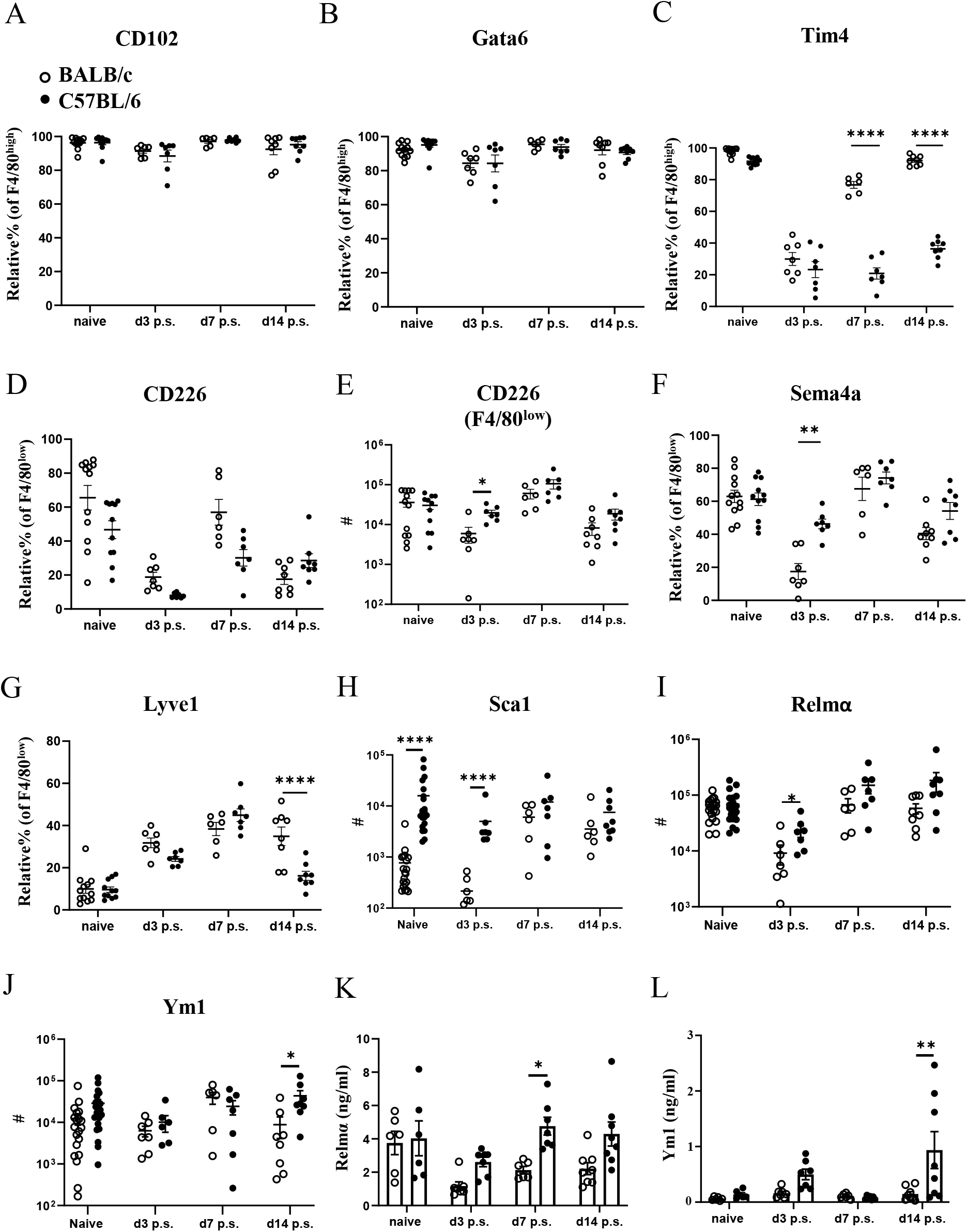
Peritoneal macrophage markers and cytokine profiles post-surgery in BALB/c mice were altered compared with C57BL/6 mice. Relative % of CD102+ (A), Gata6+ (B), and Tim4+ (C) of F4/80^high^ population. Relative % and total cell number of CD226+ F4/80^low^ (D&E, respectively). Relative % of Sema4a+ (F) and Lyve1+ (G) of F4/80^low^ macrophage population. Total cell number of Sca1+ (H), Relmα+ (I), and Ym1+ (J) in myeloid cells (CD11b+lin-Ly6C-cells)/total peritoneal cavity macrophages by flow cytometry. Peritoneal lavage levels for Relmα (K) and Ym1 (L) by ELISA. Data shows mean±SEM, pooled from 6 separate experiments for naïve or 2 separate experiments for day 3, 7, and 14 p.s., n= 6-20 mice/group; *P<0.05, **P<0.01, ****P<0.0001, ANOVA with Tukey’s multiple comparison test.

Abdominal surgery in C57BL/6 mice has previously been shown to induce the macrophage disappearance reaction of tissue resident F4/80^high^ macrophages (27). We also observed reduced numbers of F4/80^high^ macrophages in the peritoneal cavity 3 days after surgery in both C57BL/6 as well as BALB/c mice (Figure 1F). In addition, CD226 (aka DNAM-1) is highly expressed by F4/80^low^ macrophages present in the peritoneal cavity under steady state conditions (28) and these homeostatic CD226+ cells are displaced from the peritoneal cavity under inflammatory conditions (29) similar to the disappearance reaction described for F4/80^high^ resident macrophages (30). In line with this, the proportion of CD226+ cells within F4/80^low^ macrophages as well as the absolute number were reduced at day 3 after surgery in C57BL/6 and BALB/c mice compared with naïve animals (Figure 2D, E). Of note, C57BL/6 mice harboured significantly more CD226+ F4/80^low^ as well as F4/80^high^ macrophages at day 3 after surgery as compared with BALB/c mice (Figure 2E, Figure 1F). These data suggest that both mouse strains undergo the macrophage disappearance reaction but differences exist in the re-population dynamics. Semaphorin 4a (Sema4a) is expressed by circulating inflammatory monocytes and is upregulated on peritoneal macrophages during inflammation (31). Furthermore, it is retained during resolution of inflammation on F4/80^int^ and F4/80^high^ cells of monocytic origin (14). In line with a lower monocyte-derived macrophage influx on day 3 post-surgery, BALB/c mice showed fewer Sema4a+F4/80^low^ macrophages, whereas Sema4a expression was maintained on F4/80^low^ macrophages in C57BL/6 mice (Figure 2F). Similarly, Lymphatic Vessel Endothelial Receptor 1 (Lyve-1) expression has been associated with differentiation of monocyte-derived cells to F4/80^int^ macrophages (32). Interestingly, we found a significant decrease in Lyve-1 expression by F4/80^low^ macrophages in C57BL/6 mice compared with BALB/c mice on day 14 post-surgery (Figure 2G), but we found no difference in the proportion of F4/80^int^ or F4/80^high^ macrophages expressing Lyve-1 between the strains (supplementary Figure 7). This may indicate that following surgery, there is an enhanced progression of monocyte-derived cells to F4/80^int^ cells in C57BL/6 but not BALB/c mice. Of note, we found that none of the markers previously associated with specific macrophage subsets (e.g. CD102, Tim4, Lyve-1) were exclusively expressed by only one peritoneal macrophage subset (F4/80^low^, F4/80^int^, F4/80^high^) but were shared between several subsets (supplementary Figure 8). This indicates a certain degree of phenotypic plasticity between the described macrophage subsets rather than exclusive expression patterns.

With regard to cellular activation, we did not detect a significant induction of either M1 (Sca1, TNFα) or M2 (Relmα, Ym1, CD206) activation markers in response to surgery in either BALB/c or C57BL/6 mice (Figure 2 G-I, supplementary Figure 7). However, BALB/c mice showed a significantly lower number of Sca-1+ myeloid cells (CD11b+lin-Ly6C-cells) prior and at day 3 post-surgery (Figure 2H), as well as a lower number of Relmα+ cells at day 3 post-surgery (Figure 2I) and lower numbers of Ym1+ cells at day 14 after surgery compared with C57BL/6 (Figure 2J). We also analysed cytokine levels in the peritoneal lavage of BALB/c and C57BL/6 mice prior and after surgery (Figure 2K, L, supplementary Figure 9). Interestingly, the level of Relmα in the peritoneal lavage of BALB/c mice was significantly lower compared with C57BL/6 at day 7 post-surgery (Figure 2K), a slight delay compared with the Relmα+ cell reduction found at day 3 post-surgery with flow cytometry (Figure 2I). Similar to the flow cytometry data (Figure 2J), a significantly lower concentration of Ym-1 in the peritoneal lavage of BALB/c mice was detectable at day 14 post-surgery compared with C57BL/6 (Figure 2L). The remaining cytokines did not show a significant difference in levels between BALB/c and C57BL/6 mice prior and post-surgery (supplementary Figure 9). These data indicate that in addition to a reduced early influx of monocyte-derived macrophages in BALB/c mice, these cells also failed to fully integrate into the resident macrophage pool during the later resolution phase of repair following surgery.

### Genetically altered mice lacking circulating monocytes develop significantly more adhesions

Our findings indicated that BALB/c mice were more likely to have increased peritoneal adhesions at day 14 after surgery, compared with C57BL/6 mice, potentially due to reduced monocyte infiltration early following surgery. Therefore, adhesion-inducing surgery was performed in *Ccr2-* deficient mice (*Ccr2*KO on a C57BL/6 background). Due to the targeted disruption of the *Ccr2* gene, *Ccr2*KO mice fail to recruit monocytes to sites of inflammation/injury, in part because of defective egress from the bone marrow (33). As expected, following surgery, *Ccr2*KO mice showed a significantly higher number of peritoneal adhesions as well as adhesion score at day 7 compared with WT littermates (Figure 3A, B, C). *Ccr2*KO mice also showed a significantly higher relative percentage of F4/80^high^ macrophages after surgery (Figure 3D), but there was no significant difference in the total number of F4/80^high^ resident macrophages between *Ccr2*KO and WT littermates prior and post-surgery (Figure 3E). Thus, it seems unlikely that the difference in adhesion formation was due to changes in the dynamics of F4/80^high^ resident macrophages. In contrast, and in line with reduced monocyte infiltration, the relative percentages and total numbers of infiltrating monocytes (Figure 3F, G), F4/80^low^ monocyte-derived macrophages (Figure 3H, I), and F4/80^int^ converting cells (Figure 3J, K) were significantly lower in *Ccr2*KO mice compared with WT littermates after surgery. The data thus supports the hypothesis that infiltrating monocyte-derived cells play a protective role in surgery-induced peritoneal adhesion formation.

**Figure 3.**
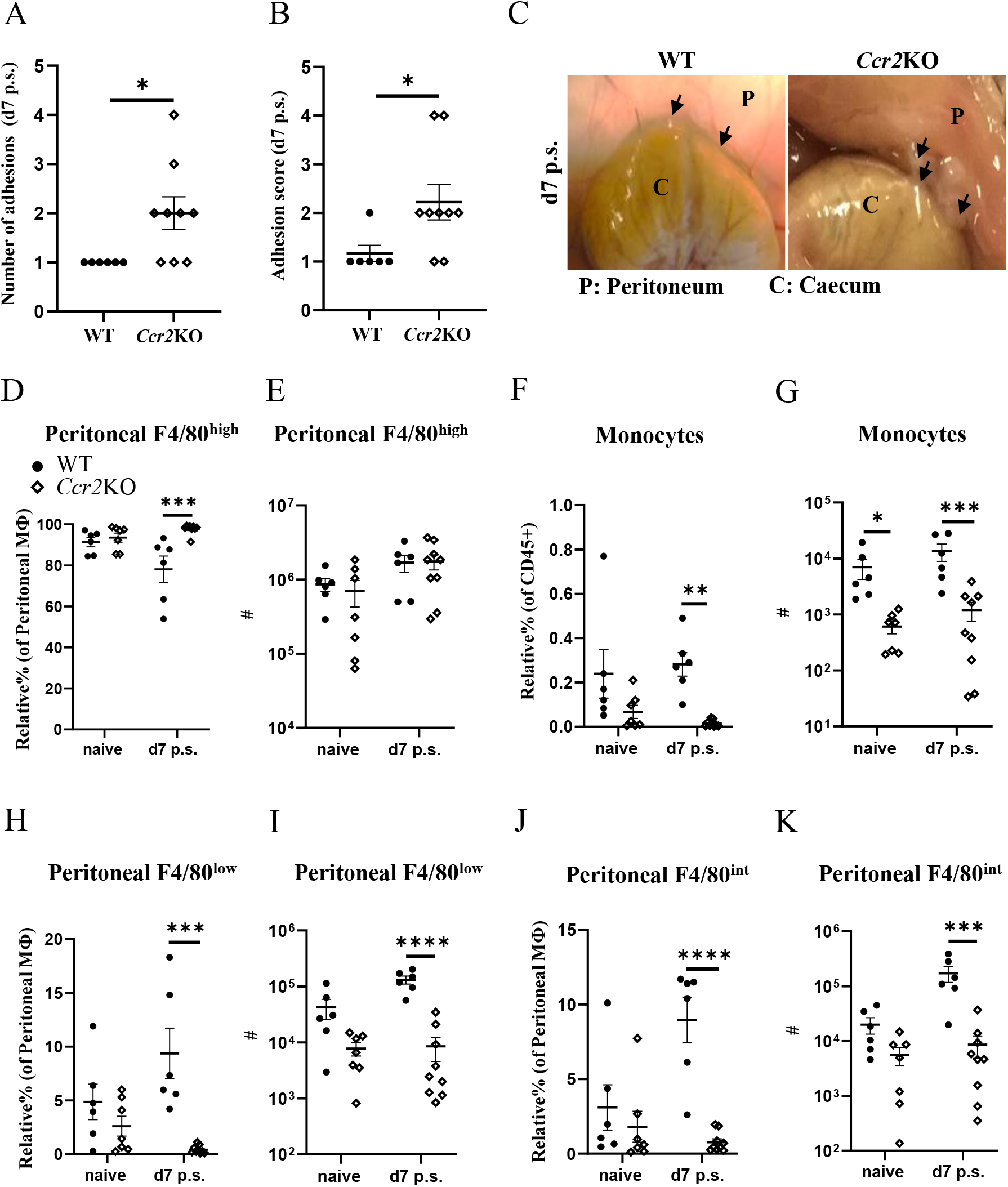
*Ccr2-*deficient mice were more susceptible to peritoneal adhesion formation post-surgery compared with their WT littermates. Number of adhesions (A), adhesion scores (B) and representative adhesion image at day 7 after surgery (C).Relative percentage and total cell number of peritoneal F4/80^high^ resident macrophages (D&E, respectively)), monocytes (F&G, respectively), peritoneal F4/80^low^ monocyte-derived macrophages (H&I, respectively), and peritoneal F4/80^int^ macrophages (J&K, respectively). Data shows mean±SEM, pooled from 5 separate experiments for naïve or 4 separate experiments for day 7, n= 6-9 mice/group; mixed male and female; *P<0.05, ***P<0.001, ****P<0.0001; (A and B) Mann Whitney test, (D-K) ANOVA with Tukey’s multiple comparison test.

### Monocyte-to-F4/80^high^ macrophage conversion was abrogated in *Ccr2*KO mice

As expected, the lack of *Ccr2* expression in *Ccr2*KO mice reduced monocyte influx into the peritoneal cavity compared with WT littermates. Consequently, similar to our data with BALB/c mice (Figure 2C), the relative percentage of Tim4+ cells within F4/80^high^ macrophages at day 7 after surgery in *Ccr2*KO mice was significantly higher compared with WT littermates (Figure 4A). The lack of monocyte influx in *Ccr2*KO mice also caused significantly lower relative percentage of Sema4a+ cells within F4/80^low^ (Figure 4B), F4/80^int^ (Figure 4C), and F4/80^high^ macrophage subsets (Figure 4D) compared with WT littermates. In addition, the relative percentage of Lyve-1+ cells within F4/80^low^ (Figure 4E), F4/80^int^ (Figure 4F), and F4/80^high^ (Figure 4G) macrophage subsets in *Ccr2*KO mice was also significantly lower compared with WT littermates. The relative percentage of CD226 within F4/80^low^ macrophage population did not show any significant difference between the groups (Figure 4H), but *Ccr2*KO mice had significantly lower total number of CD226+ cells compared with WT littermates (Figure 4I). We also analysed the activation of all macrophage populations and found that *Ccr2*KO peritoneal macrophages had significantly lower CD206 (Figure 4J) and Relmα expression (Figure 4K) prior and post-surgery compared with WT littermates. Similar to our flow cytometry data, the level of Relmα in the peritoneal lavage were also significantly lower in *Ccr2*KO at day 7 post-surgery compared with WT littermates (Figure 4L). The levels of other cytokines, including Ym1, IFN-γ, IL-6, IL-10, IL-12, and IL-13 remained the same between the groups (supplementary Figure 10).

**Figure 4.**
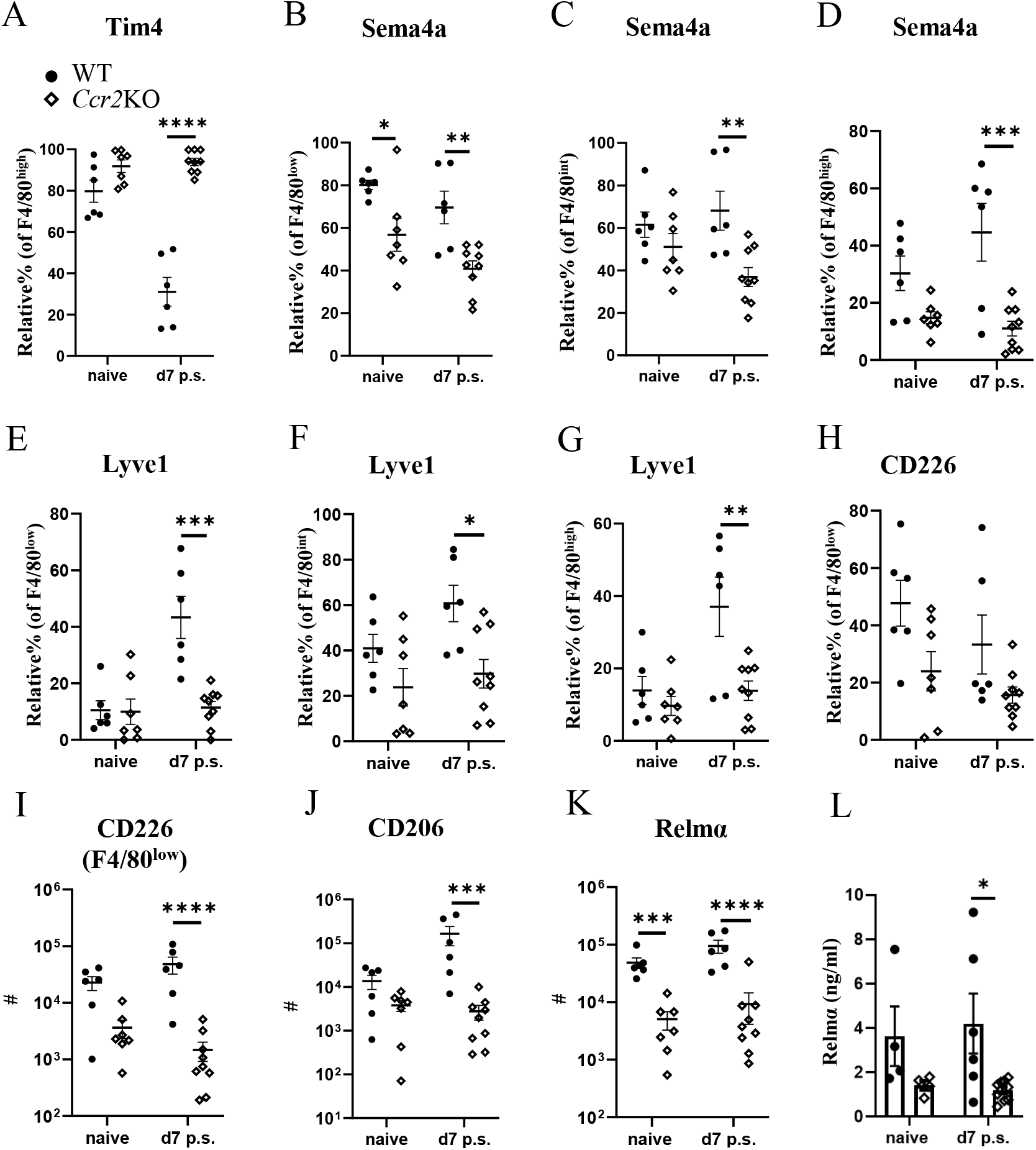
Macrophages markers associated with monocyte-derived and intermediate subsets were altered in *Ccr2-*deficient mice. Relative % of Tim+ (A) within F4/80^high^ population, Relative % of Sema4a+ within F4/80^low^ (B), F4/80^int^ (C), and F4/80^high^ (D). Relative % of Lyve1+ within F4/80^low^ (E), F4/80^int^ (F), and F4/80^high^ (G). Relative % of CD226+ within F4/80^low^ (H). Total number of CD226+ cells within F4/80^low^ subset (I) Total number of CD206+ (J) and Relmα+ (K) cells within total peritoneal macrophages. Cytokine level of Relmα in peritoneal lavage (L) using ELISA. Data shows mean±SEM, pooled from 4-5 separate experiments for naïve or 4 separate experiments for day 7, n= 4-6 mice/group; mixed male and female, *P<0.05, **P<0.01, ***P<0.001, ****P<0.0001, ANOVA with Tukey’s multiple comparison test.

### Depletion of monocyte-derived peritoneal macrophages induced more peritoneal adhesions post-surgery

As *Ccr2*KO mice do not express the *Ccr2* gene, monocyte influx is defective throughout their life course, thus, it is not possible to determine whether early influx of monocytes, as observed in C57BL/6 mice on day 3 post-surgery (Figure 1G), was contributing to the reduction in adhesions or whether influx of monocytes at a later stage (day 7 or 14 post-surgery) was also of importance. In addition, due to effects on their replenishment, phenotypically distinct resident macrophages accumulate in *Ccr2*KO animals with age (26). Therefore, the observed susceptibility to adhesion formation may have been due to altered functional responses of cavity resident macrophages in *Ccr2*KO mice rather than defective monocyte recruitment. To address this issue, C57BL/6 mice were treated with an anti-CCR2 monoclonal antibody (mAb; clone MC21; (23) injected i.p. at day - 1, 0, and 1 after surgery) to specifically deplete monocytes during the early phases of repair. Rat IgG2b was used as isotype control, and mice were analysed at day 14 post-surgery (Figure 5A). MC21 mAb has been shown to effectively deplete CCR2+ (CD11b+Ly6C^high^) monocytes (23). Similar to our findings with BALB/c and *Ccr2*KO mice, early depletion of infiltrating monocytes enhanced the susceptibility of C57BL/6 mice to surgery-induced adhesions at day 14 post-surgery compared with isotype control-antibody treated mice (Figure 5B, C, D). Moreover, the site-specific adhesion tissue in anti-CCR2 mAb treated mice also showed significantly higher collagen deposition at day 14 post-surgery compared with isotype control treated mice (Figure 5D, E). Both at day 3 as well as day 14 post-surgery, no significant difference in the total number of PECs (Figure 5F), F4/80^high^ macrophages (Figure 5G) or monocytes (Figure 5H) could be detected between the groups. However, anti-CCR2 mAb treated mice regained significantly higher Tim4 expression within F4/80^high^ macrophage subset at day 14 post-surgery compared with isotype control-treated mice (Figure 5I), indicative of reduced monocyte-integration in anti-CCR2 Ab treated mice. Of note, unlike BALB/c (Figure 2C) and *Ccr2*KO mice (Figure 4A), anti-CCR2 mAb treated mice only partially regained Tim4 expression (∼ 44%) and did not reach the same level as detected in naïve C57BL/6 mice (∼ 92%, Figure 2C), suggesting a delayed integration of monocyte-derived cells. In line with this hypothesis, no significant difference in any of the other peritoneal macrophage markers (e.g. Sema4a, Lyve-1) was found between anti-CCR2 mAb treated and isotype control-treated mice after surgery (supplementary Figure 11 and 12). Interestingly, the level of Relmα in the peritoneal lavage of anti-CCR2 mAb-treated mice at day 14 post-surgery was significantly higher compared with isotype control-treated mice (Figure 5J), which may indicate a delay in the induction of surgery-induced responses (Figure 2K) or over-compensatory mechanisms. All other cytokines remained similar between the groups (supplementary Figure 13). Together, the treatment of C57BL/6 mice with a monocyte-depleting antibody thus provided direct evidence that monocytes contribute to protection against adhesion formation.

**Figure 5.**
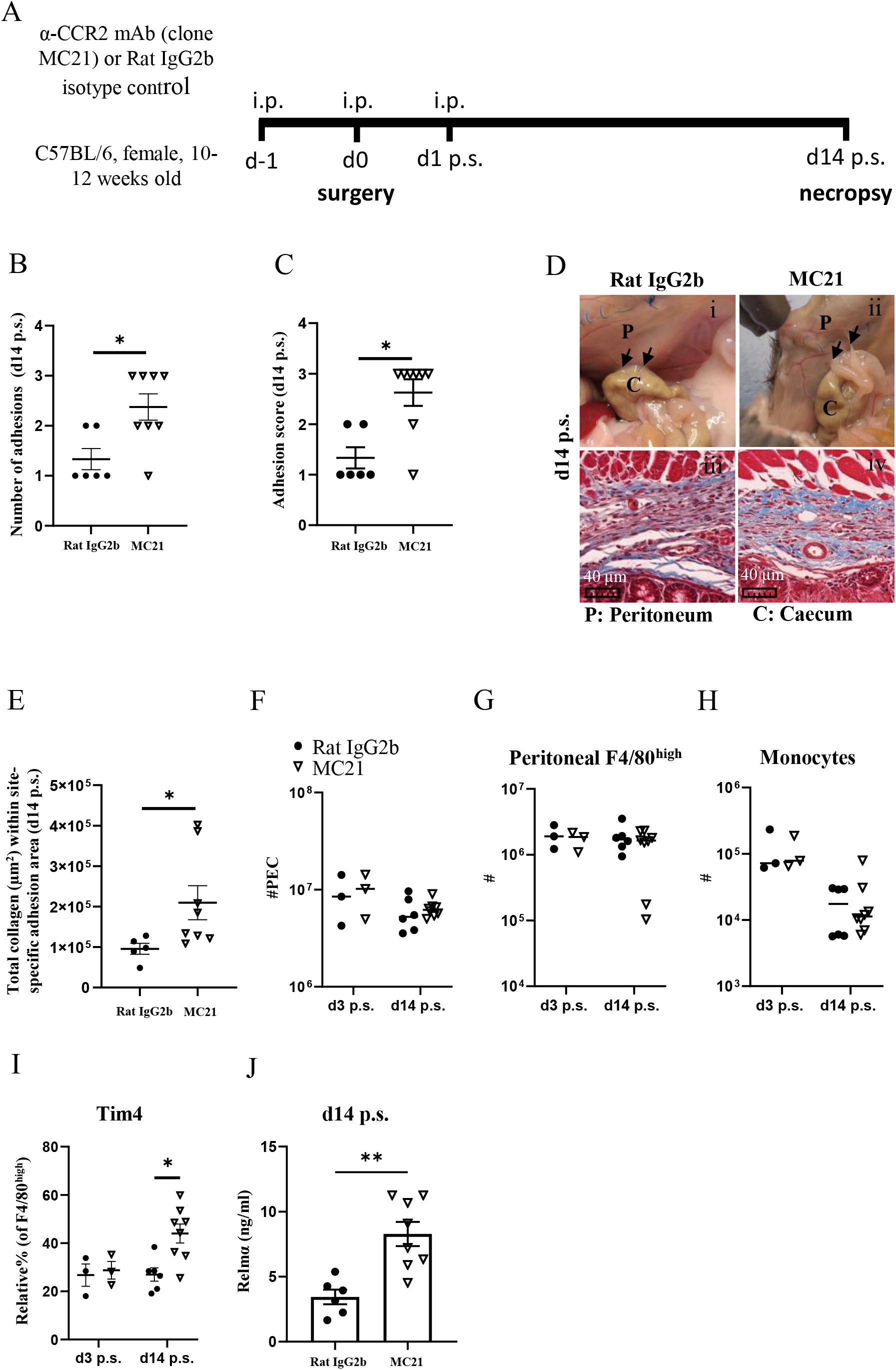
Depletion of monocyte-derived peritoneal macrophages enhanced susceptibility of C57BL/6 mice to peritoneal adhesions post-surgery. Schematic of experiment approach (A). Number of adhesions (B) and adhesion score at day 14 post-surgery (C). Representative image and corresponding histological appearance (Masson’s trichrome staining) of adhesions at day 14 in mice treated with CCR2-mAb or control IgG2b (D). Collagen analysis in site-specific adhesion histological sections at day 14 after surgery(E). Total peritoneal exudate cells (F). Total cell number of peritoneal F4/80^high^ resident macrophages (G) and monocytes (H) at day 3 and 14 post-surgery. Relative % of Tim4+ cells within F4/80^high^ population (I). Level of Relmα in peritoneal lavage at day 14 (J). Data shows mean±SEM, pooled from 2 separate experiments, n= 6-8 mice/group; *P<0.05, **P<0.01; (B, C, E J) Mann Whitney test, (F-I) ANOVA with Tukey’s multiple comparison test. Scale bar 40μm.

## Discussion

The contribution of peritoneal macrophages to adhesion formation after surgery remains ill-defined, with both a protective and pathological role reported (9, 11, 12). Furthermore, mouse strain differences in susceptibility to surgical adhesion formation following pneumoperitoneum/laparoscopy and uterine horn/side wall trauma was demonstrated in previous study (19), but the mechanisms driving these differences remain largely unexplored. Here, we found that C57BL/6 mice showed a significantly lower peritoneal adhesion score after surgery compared with BALB/c mice in agreement with the previous study (19). However, unlike that previous study (19), we found that the susceptibility of BALB/c mice to peritoneal adhesion did not become apparent until day 14 post-surgery. Moreover, site-specific adhesions that did form displayed a relatively lower collagen deposition suggesting the adhesions maybe less mature and more readily degraded. Importantly, C57BL/6 mice showed an enhanced influx of monocytes and monocyte-derived macrophages at day 3 post-surgery. Blocking the recruitment of circulating monocytes in C57BL/6 mice, either by using *Ccr2*KO mice or antibody-mediated CCR2 depletion, resulted in an enhanced susceptibility to peritoneal adhesions after surgery. Together, these data provide evidence that circulating monocytes or monocyte-derived macrophages help prevent peritoneal adhesions following surgery (Figure 6).

**Figure 6.**
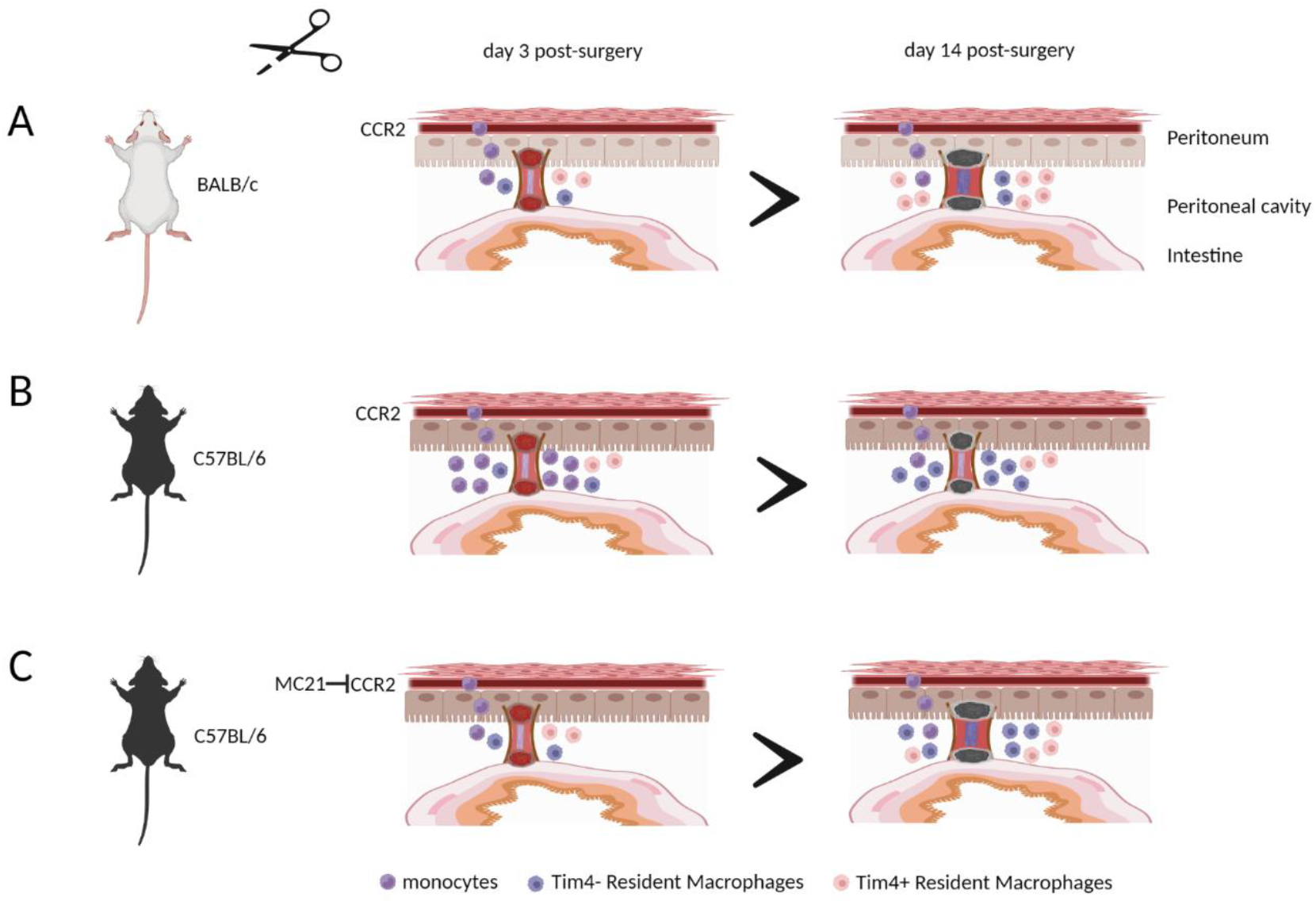
Monocytes and/or monocyte-derived peritoneal macrophages are essential in preventing peritoneal adhesion formation after surgery. BALB/c mice showed a significantly lower influx of monocytes and monocyte-derived macrophages at day 3 post-surgery, leading to greater peritoneal adhesion formation at day 14 post-surgery (A) compared with C57BL/6 mice (B). Blocking the recruitment of circulating monocytes in C57BL/6 mice resulted in an enhanced susceptibility to peritoneal adhesions after surgery (C). The illustration was created using Biorender.com.

Our data complements and advances two previous studies of surgery-induced adhesion formation, in which experimental enhancement of monocyte recruitment showed a protective effective. In a rabbit study, recruitment of inflammatory cells, including monocytes, by intraperitoneal injection of protease peptone prior to surgery reduced the number of peritoneal adhesions formed (9). In a mouse model of ischaemic button-induced adhesion formation, thioglycolate or monocyte chemoattractant protein-1 (MCP-1) injection increased inflammatory monocyte influx and reduced adhesion formation (11). In this latter study, transient neutrophil depletion was also proposed as a complementary mechanism for reducing adhesions (11). In this context, it is interesting to note, that although BALB/c mice were more susceptible to peritoneal adhesions in our model, an equivalent increase in neutrophils in the peritoneal cavity at day 3 after surgery was found compared with C57BL/6 mice (supplementary Figure 14). The relative percentage and total cell number of other population, including eosinophils, B cells, and T cells are also shown in supplementary Figure 14. Moreover, *Ccr2*KO mice have been shown to recruit equivalent numbers of neutrophils in a model of bacteria-induced inflammation (34). Thus, the protective effect of monocytes or monocyte-derived cells in our model seems unlikely to be dependent on an altered neutrophil response.

Similarly, two recent studies suggested that rapid recruitment of F4/80^high^ peritoneal resident macrophages to the site of mesothelial injury may result in the creation of a barrier preventing excessive fibrin attachment and consequently preventing the formation of adhesions after surgery (25, 35). We found that C57BL/6 mice showed a significantly higher number of peritoneal F4/80^high^ resident macrophages prior and at day 3 post-surgery compared with BALB/c mice. Thus, C57BL/6 mice may be inherently protected from peritoneal adhesion formation by forming a macrophage-rich barrier. However, the number and dynamics of the resident macrophage population in both *Ccr2*KO and antibody-depleted mice were not altered compared with their respective controls, while these mice were still more susceptible to peritoneal adhesion formation. Our data from *Ccr2*KO and anti-CCR2 mAb depleted animals, both on a C57BL/6 background, therefore argue for an additional protective role for monocytes and monocyte-derived cells as described, independent of a possible protective barrier function of F4/80^high^ resident macrophages.

Of note, we found that the difference in adhesion score between BALB/c and C57BL/6 mice occurred at a later stage of adhesion formation (day 14 post-surgery). Whilst the adhesion score in BALB/c mice remained the same between day 7 and day 14 after surgery, C57BL/6 mice showed a reduced adhesion score at day 14 suggesting that some adhesions that initially formed in C57BL/6 may have been immature and cleared possibly due to lower collagen deposition to stabilise development. Clearance of early fibrin-rich adhesions has been associated with increased fibrinolysis in the initial stages post-surgery (4) and so the protective effect of monocytes or monocyte-derived macrophages may be due to promotion of proteolytic activity and hence degradation of collagen-poor unstable adhesions between days 7 and 14 post-surgery. This proposal hypothesis is consistent with the finding that monocyte-derived macrophages degrade fibrin-rich aggregates that form in response to intraperitoneal bacterial infection (36). Interestingly, the response of resident F4/80^high^ Gata6 dependent macrophages was found to depend on the severity of the insult (35). Minor injury (laser injury) induced cavity resident macrophages to aggregate on the injured site to form a temporary wound covering or barrier. However, with more severe injury (ischaemic button), these aggregates expanded to form superaggregates spanning adjacent tissues and became a nidus for subsequent adhesion formation (35). Thus, the protective effects of monocytes or monocyte-derived macrophages in our model may be explained by active degradation of immature adhesions through fibrinolytic activity or by disrupting F4/80^high^ resident macrophage-derived superaggregates.

Importantly, we cannot distinguish whether the protective effect is mediated via monocytes themselves or via macrophages that have differentiated from monocytes in the cavity (Bain et al., 2016). Differences in the recruitment of monocytes to the peritoneal cavity between BALB/c and C57BL/6 mice were largely restricted to the early phases of the repair process (day 3 post-surgery, Figure 1F) and early but transient depletion of monocytes significantly affected adhesion formation (Figure 5B). Changes in monocyte-derived macrophage populations (F4/80^low^ and F4/80^int^) were slightly prolonged, but similarly temporary (Figure 1G, H). Thus, it is unclear if these cells act early during the adhesion formation process, but the outcome of these actions was not visible until later (day 14 post-surgery), or whether the cells acquire an activation phenotype at later stages dependent on their earlier influx. Interestingly, we found a marked change in the phenotype of F4/80^high^ resident macrophages at later stages as indicated by the loss of Tim4 expression by these cells (Figure 2C). Tim4 has been linked to the embryonic origin of resident macrophages, while a loss of Tim4 expression indicates integration of monocyte-derived macrophages into the resident pool (24). Thus, the maintained expression of Tim4 by F4/80^high^ macrophages in BALB/c, *Ccr2*KO, and partially in anti-CCR2 mAb-treated mice indicates a failure to integrate monocyte-derived cells into the resident pool. Importantly, these monocyte-derived resident cells are functionally different to their embryonic-derived counterparts (14). Therefore, it is possible that the protective effect of monocytes is exerted at the level of monocyte-derived, F4/80^high^ Tim4-ve resident cells. In this case, the reason why we do not observe notable differences during the early stages of the adhesion process may be due to the delay necessitated by the differentiation process of these cells.

Overall, our data suggest that several macrophage populations possess the capacity to prevent adhesion formation, with each acting at different phases of the resolution process (eg. resident cells early versus. monocyte-derived cells later) and utilising different, non-redundant mechanisms (eg. barrier-function versus fibrinolysis), respectively. Importantly, the effectiveness of these mechanisms is likely to be further influenced by other factors like severity of the injury (35), the presence of inflammatory stimuli (37) or neutrophil infiltration (11) as well as a possible genetic differences as shown here. Our findings reveal an important role of monocyte-derived cells to prevent adhesion formation during the recovery phase after surgery. The results of our current study may inform the idea that manipulation of monocytes/macrophages may in future be used to ameliorate surgical adhesions. More work, however, is needed in experimental models before this can be translated to humans. It has previously been suggested that injection of an artificial CCR2 ligand (recombinant MCP-1) to attract monocytes to the peritoneal cavity at the time of surgery can reduce adhesion formation in mice, however the effect did not reach statistical significance (11) and MCP-1/CCL2 is not specific for monocytes but also attracts memory T lymphocytes and natural killer (NK) cells (38). Thus, such a treatment may yield unwanted side-effects. Other chemokines acting via CCR2 (e.g. PC3-secreted microprotein (PSPM)) have been suggested to more specifically target monocytes, but their effect on adhesion formation is not known (39). In addition, chemokines or any CCR-2 based treatment may be hampered by genetic polymorphisms in the CCR-2 gene (40).. As an alternative systemic CSF-1 delivery has been shown to increase intestinal monocyte and MHC-II^low^ macrophage numbers overcoming CCR2 deficiency and promoting the resolution of gut inflammation (41) and, thus, could be adapted for the peritoneal cavity. However, caution is needed as bacterial contamination after surgery can enhance adhesion formation in an EGFR-dependent manner and, importantly, bone marrow-/monocyte-derived macrophages are the main source of EGFR-ligands under these circumstances (37). Therefore, in contrast to our relatively sterile conditions, monocytes/monocyte-derived cells may actually enhance adhesion formation in the presence of bacterial contaminants. Thus, further research is required to identify conclusively the specific phenotype of monocyte-derived cells harbouring this protective capacity as well as to decipher their pivotal mechanism of action. Such data will allow the development of an innovative approach in preventing peritoneal adhesion maturation by modulating monocyte-derived macrophages following surgery.

## Data Availability Statement

All data sets supporting this study are available on request.

## Ethics statement

All animal experiments were approved by the University of Manchester Animal Welfare and Ethical Review Board and performed under the regulation of the Home Office Scientific Procedures Act (1986) and the Home Office approved licence (P1208AD89).

## Author Contributions

RS: designed and performed experiments, analyzed data, and wrote the manuscript. KD: perfomed the surgery and edited the manuscript. DR and SH: designed experiments, analyzed data, wrote the manuscript and obtained funding. ASW and JA: designed experiments, provided input for interpretation, edited the manuscript and obtained funding. MM: provided input for interpretation and edited the manuscript.

## Acknowledgements

We would like to thank John Grainger and Kara Filbey for sharing the *Ccr2*-deficient mice. We would also like to thank Gareth Howell from the flow cytometry facility, staff in the Bioimaging facility and BSF facility at the University of Manchester, and Stella Pearson for technical assistence sectioning histology samples. Some aspects of this manuscript have previously been released as a preprint (42).

## List of Supplementary Material

Supplementary Table 1. Adhesion scoring system

Supplementary Table 2. Panel 1 flow cytometry

Supplementary Table 3. Panel 2 flow cytometry

Supplementary Table 4. The coating and detection antibody for ELISA

Supplementary Figure 1. Graphical of representation of the experimental approach to induce adhesion formation after surgery.

Supplementary Figure 2. Gating strategy for flow cytometry analysis of PECs with the representative of lineage+ lineage- and F4/80 against IA/IE plots between BALB/c and C57BL/6 mice prior and after surgery.

Supplementary Figure 3. Gating strategy for flow cytometry analysis of lineage+ population.

Supplementary Figure 4. Exemplary illustration on how total collagen on site-specific adhesion area was measured.

Supplementary Figure 5. Representative of adhesion images and Masson’s trichrome staining and total collagen at day 3 and 7 post-surgery between BALB/c and C57BL/6 mice.

Supplementary Figure 6. Peritoneal exudate cells and relative percentage of peritoneal macrophages and monocytes prior and after surgery.

Supplementary Figure 7. Relative percentages for different markers expressed by peritoneal macrophages within BALB/c and C57BL/6 mice prior and post-surgery.

Supplementary Figure 8. Schematic representation of how different markers of peritoneal macrophages were expressed in different subsets.

Supplementary Figure 9. Cytokine analysis in the peritoneal lavage of BALB/c and C57BL/6 before and after surgery.

Supplementary Figure 10. Cytokine analysis in the peritoneal lavage of Ccr2KO and WT littermate before and after surgery.

Supplementary Figure 11. Relative percentage of monocytes and different markers of peritoneal macrophages of Rat IgG2b isotype control and anti-CCR2 (clone MC21) treated mice after surgery.

Supplementary Figure 12. Total cell number of monocytes and different markers of peritoneal macrophages of Rat IgG2b isotype control and anti-CCR2 (clone MC21) antibody treated mice after surgery.

Supplementary Figure 13. Cytokine analysis in the peritoneal lavage of Rat IgG2b isotype control and anti-CCR2 (clone MC21) antibody treated mice after surgery.

Supplementary Figure 14. The relative percentage and total cell number of neutrophils, eosinophils, B cells, and T cells in the peritoneal cavity of BALB/c and C57BL/6 mice prior and after surgery.

