## supplementary for "Monocyte-derived peritoneal macrophages protect C57BL/6 mice against surgery-induced adhesions"

**Supplementary Table 1. Adhesion scoring system based on the type of adhesion**

| | | **Injured peritoneum** | | **Incision site** | |
|---|---|---|---|---|---|
| | No adhesion | Caecum to injured peritoneum | Pelvic fat/omentum to injured peritoneum/caecum | Pelvic fat/omentum to incision site | Organ(s) to incision site/injury site |
| Score | 0 | 1 | 2 | 3 | 4 |

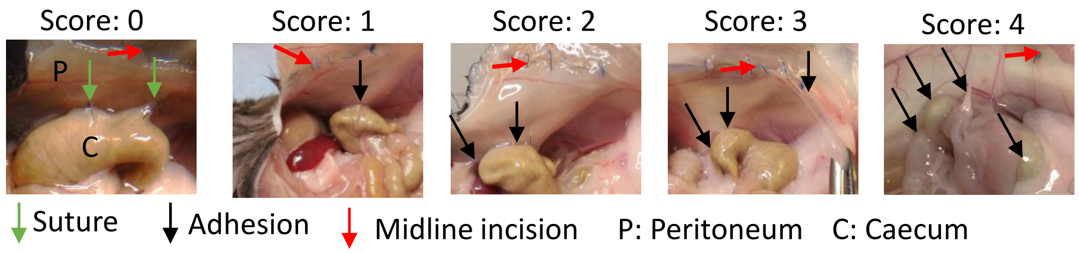

**Supplementary Table 2. Panel 1 flow cytometry**

| Filter | Fluorochrome | Antigen | Clone | Dilution | Supplier |
|---|---|---|---|---|---|
| 355 450_50 | BUV440 | Live/Dead | Zombie UV | 1:1000 | Biolegend |
| 355 735_30 | BUV737 | Ym1 biotin | BAF2446 | 1:83 | R& D System |
| 405 450_50 | PB | CD19 | 6D5 | 1:300 | Biolegend |
| 405 450_50 | BV421 | SigF | S17007L | 1:300 | Biolegend |
| 405 450_50 | PB | TCRβ | H57-597 | 1:300 | Biolegend |
| 405 450_50 | PB | Ly6G | 1A8 | 1:300 | Biolegend |
| 405 450_50 | BV421 | NK1.1 | PK136 | 1:300 | Biolegend |
| 405 450_50 | BV421 | Ter119 | TER-119 | 1:300 | Biolegend |
| 405 525_50 | BV510 | Ly6C | HK1.4 | 1:300 | Biolegend |
| 405 605_40 | BV605 | CD11c | N418 | 1:300 | Biolegend |
| 405 677_20 | BV650 | CD102 | 3C4(mIC2/4) | 1:300 | BD Biosciences |
| 405 710_50 | BV711 | CD11b | M1/70 | 1:500 | Biolegend |
| 405 750_30 | BV750-P | TNFa | MP6-XT22 | 1:166 | Biolegend |
| 405 810_40 | BV786 | Sca-1 | D7 | 1:500 | Biolegend |
| 488 530_30 | AF488 | Relm-a | 500-P214-100 | 1:166 | Peprotech |
| 488 710_50 | PerCPeF 710 | CD73 | eBioTY/11.8 | 1:833 | Invitrogen |
| 561 586_15 | PE | GATA6 | D61E4 | 1:166 | Cell Signaling Technology |
| 561 610_20 | PE-CF594 | F4/80 | BM8 | 1:300 | Biolegend |
| 561 670_30 | PE-Cy5 | CD45 | 30-F11 | 1:666 | Biolegend |
| 561 780_60 | PE-Cy7 | Tim4 | RMT4-54 | 1:833 | Biolegend |
| 637 670_30 | AF647 | CD115 | AFS98 | 1:300 | Biolegend |
| 637 730_45 | AF700 | I-A/I-E | M5/114.15.2 | 1:833 | Biolegend |
| 637 780_60 | APC-cy7 | CD226 | 10E5 | 1:300 | Biolegend |

**Supplementary Table 3. Panel 2 flow cytometry**

| Filter | Fluorochrome | Antigen | Clone | Dilution | Suplier |
|---|---|---|---|---|---|
| 355 450_50 | BUV440 | Live/Dead | Zombie UV | 1:1000 | Biolegend |
| 355 735_30 | BUV737 | CD206 biotin | C068C2 | 1:166 | Biolegend |
| 405 450_50 | PB | CD19 | 6D5 | 1:300 | Biolegend |
| 405 450_50 | BV421 | SigF | S17007L | 1:300 | Biolegend |
| 405 450_50 | PB | TCRβ | H57-597 | 1:300 | Biolegend |
| 405 450_50 | PB | Ly6G | 1A8 | 1:300 | Biolegend |
| 405 450_50 | BV421 | NK1.1 | PK136 | 1:300 | Biolegend |
| 405 450_50 | BV421 | Ter119 | TER-119 | 1:300 | Biolegend |
| 405 525_50 | BV510 | Ly6C | HK1.4 | 1:300 | Biolegend |
| 405 605_40 | BV605 | CD11c | N418 | 1:300 | Biolegend |
| 405 677_20 | BV650 | CD102 | 3C4(mIC2/4) | 1:300 | BD Biosciences |
| 405 710_50 | BV711 | CD11b | M1/70 | 1:500 | Biolegend |
| 405 810_40 | BV786 | Ki76 | B56 | 1:50 | BD Biosciences |
| 488 530_30 | AF488 | Relma | 500-P214-100 | 1:166 | Peprotech |
| 488 710_50 | PerCPeF 710 | CD73 | eBioTY/11.8 | 1:833 | Invitrogen |
| 561 586_15 | PE | Lyve-1 | 10/FR2 | 1:166 | Biolegend |
| 561 610_20 | PE-CF594 | F4/80 | BM8 | 1:300 | Biolegend |
| 561 670_30 | PE-Cy5 | CD45 | 30-F11 | 1:666 | Biolegend |
| 561 780_60 | PE-Cy7 | Tim4 | RMT4-54 | 1:833 | Biolegend |
| 637 670_30 | AF647 | sema4a | 5E3/SEMA4A | 1:166 | Biolegend |
| 637 730_45 | AF700 | I-A/I-E | M5/114.15.2 | 1:833 | Biolegend |
| 637 780_60 | APC-cy7 | CD115 | AFS98 | 1:300 | Biolegend |

**Supplementary Table 4. The coating and detection antibody for ELISA**

| Cytokine | Coating antibody clone | Supplier | Detection antibody clone | Supplier |
|---|---|---|---|---|
| IL-4 | 11B11 | Biolegend | BVD6-24G2 | Biolegend |
| IL-10 | JES5-16E3 | Biolegend | JES5-2A5 | Biolegend |
| IL-13 | eBio13A | Invitrogen | eBio1316H | Invitrogen |
| Relm-α | 500-P214 | Peprotech | 500-P214Bt | Peprotech |
| Ym1 | Duoset | R&D System | Duoset | R&D System |
| IFN-γ | Cat. 551216 | BD Biosciences | XMG1.2 | Biolegend |
| IL-12p40 | C15.6 | Biolegend | C17.8 | Biolegend |
| IL-6 | Cat. 554400 | BD Biosciences | MP5-3C11 | Biolegend |
| TNF-α | MP6-XT22 | Biolegend | MP6-XT22 | Biolegend |

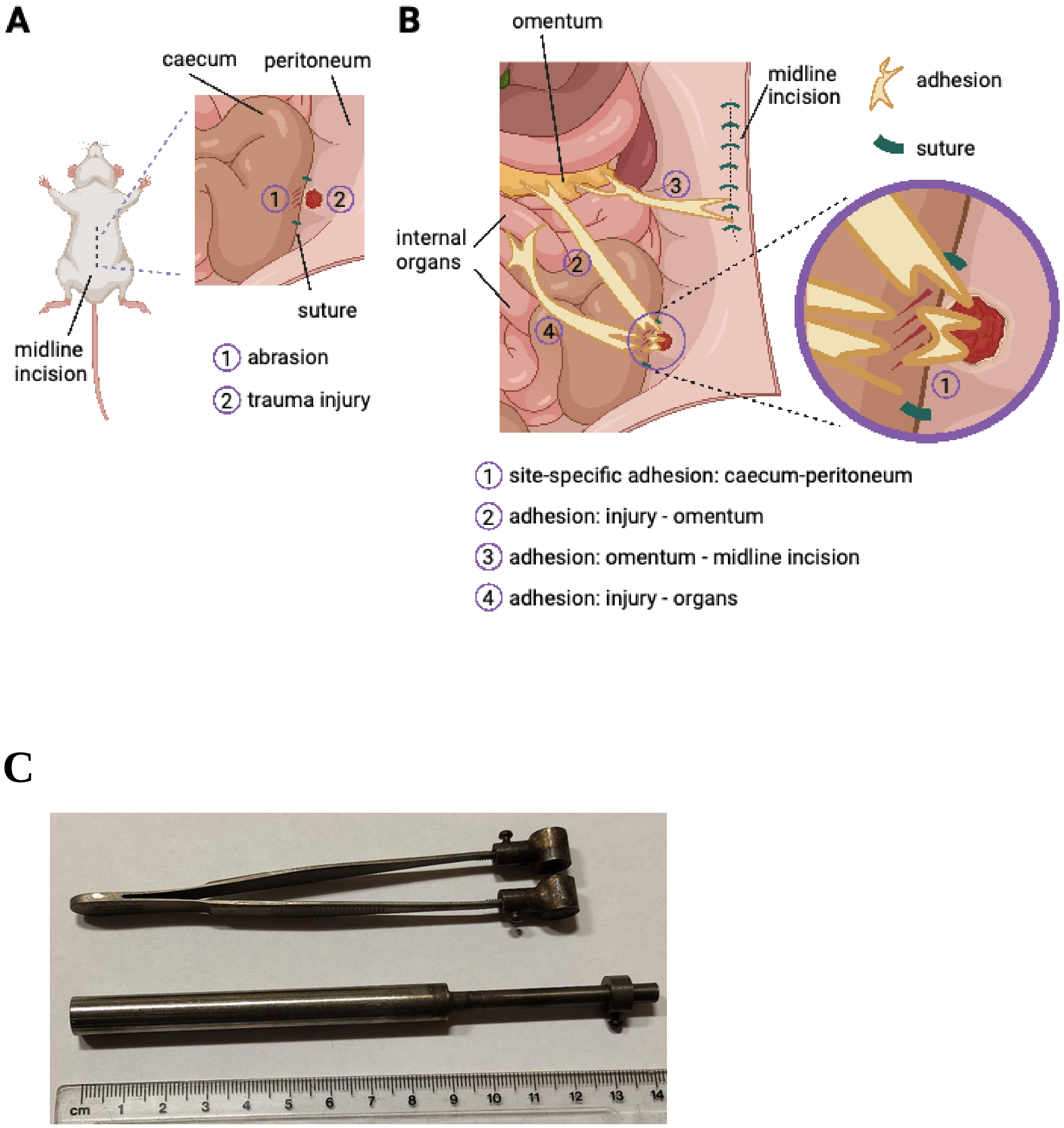


**Supplementary Figure 1.** Graphical illustration (A and B) of the experimental approach and image (C) of the trauma instrument used to induce peritoneal injury. The diagrams in A and B were created with BioRender.com.

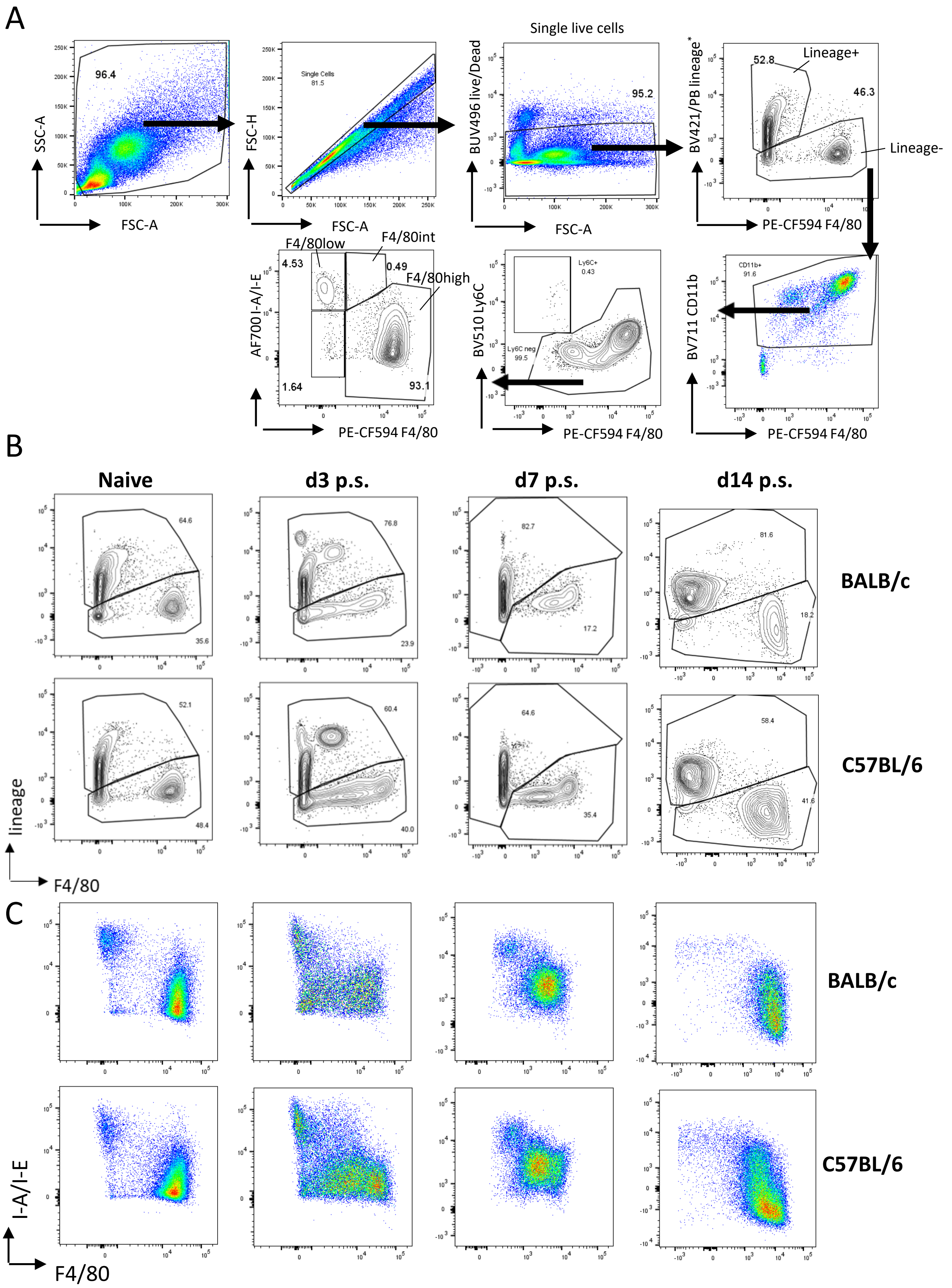


**Supplementary Figure 2. (A)** Gating strategy for flow cytometry analysis and representative gating of lineage+ or lineage- cells followed by the identification of F4/80 high, F4/80 int and F4/80 low macrophages. **(B & C)** Exemplary plots of F4/80 against lineage (B) or I-A/I-E (C) in BALB/c and C57BL/6 prior at different timepoints after surgery. Lineage* (CD19, SigF, TCRb, Ly6G, NK1.1, and Ter119)

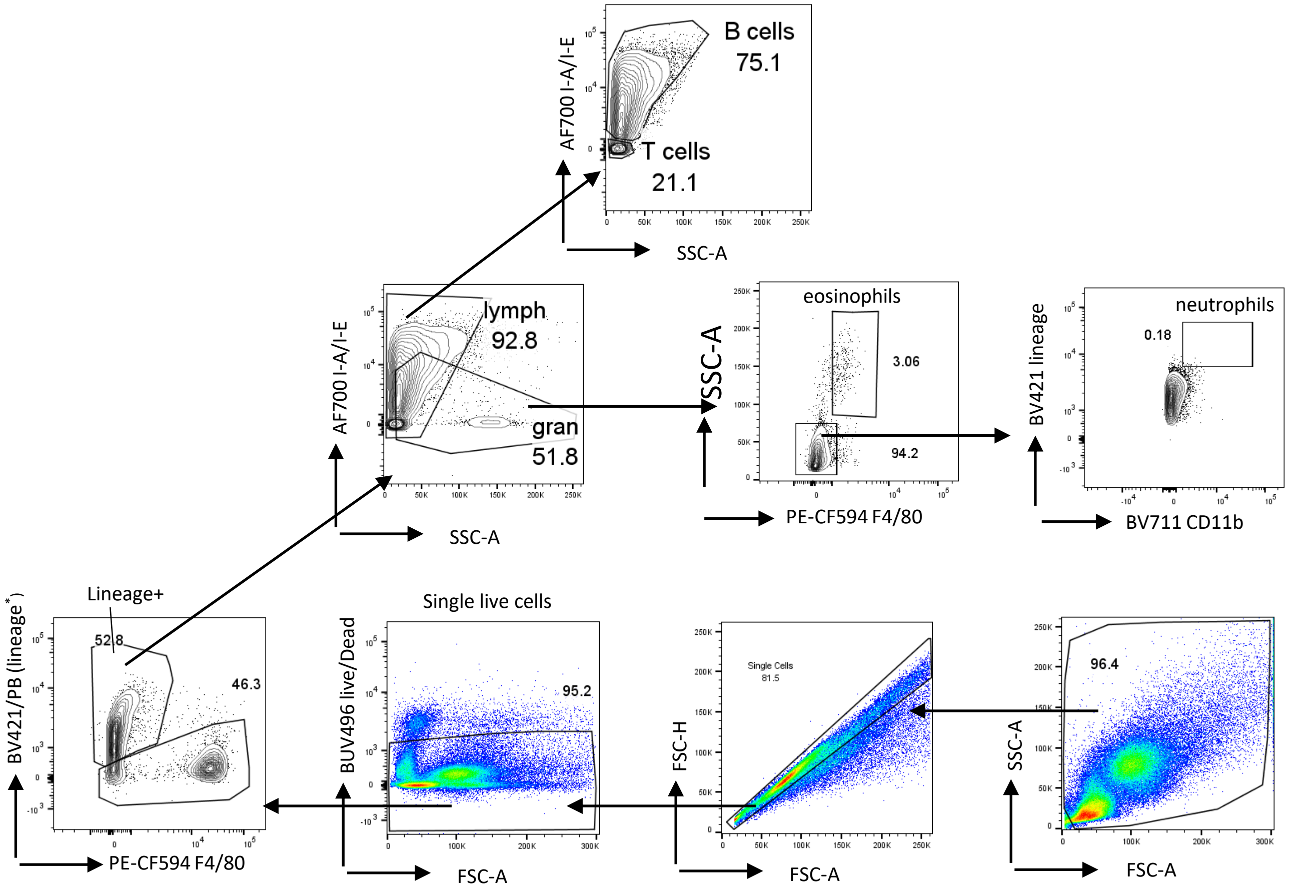


**Supplementary Figure 3.** Gating strategy for flow cytometry analysis of lineage+ populations. Lineage* (CD19, SigF, TCRb, Ly6G, NK1.1, and Ter119). lymph: lymphocytes; gran: granulocytes

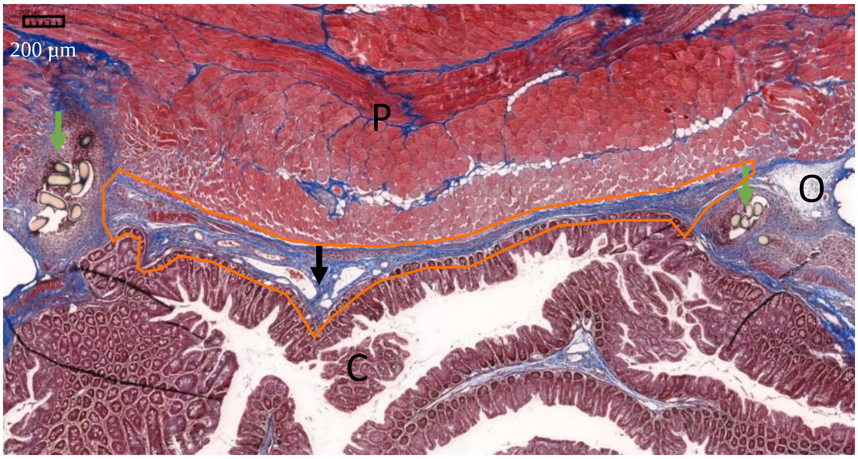


↓ Suture ↓ Adhesion P: Peritoneum C: Caecum O: Omentum

orange line: site-specific adhesion area

**Supplementary Figure 4.** Exemplary illustration on how total collagen on site-specific adhesion area was measured. Total collagen within the site specific adhesion area was measured using RGB thresholding by QuantCentre and HistoQuant plugin on SlideViewer software (version 2.5). Scale bar: 200µm

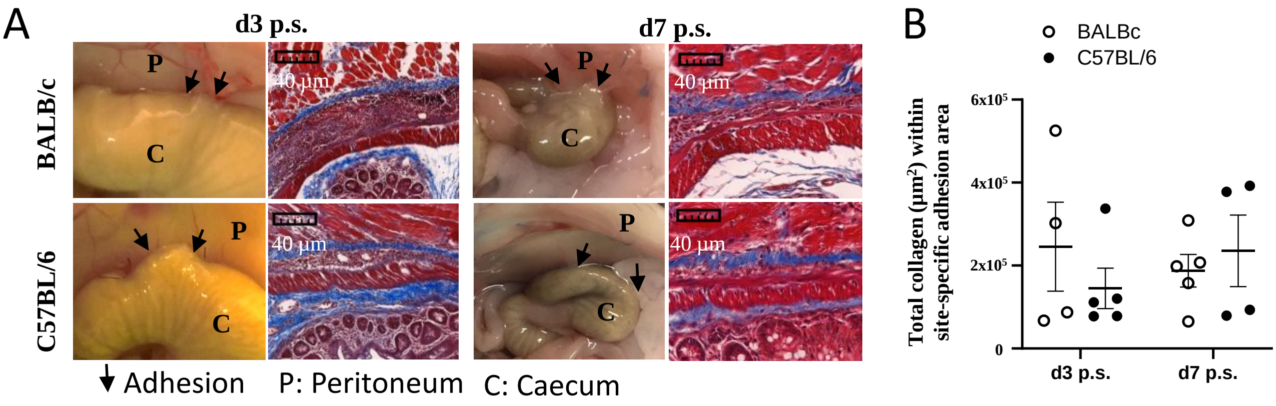


**Supplementary Figure 5. (A)** Representative photographs of adhesions and micrographs of Masson's trichrome stained histological sections at day 3 and 7 after surgery in BALB/c and C57BL/6 mice. Scale bar: 40µm. **(B)** Total collagen content in site-specific adhesions.

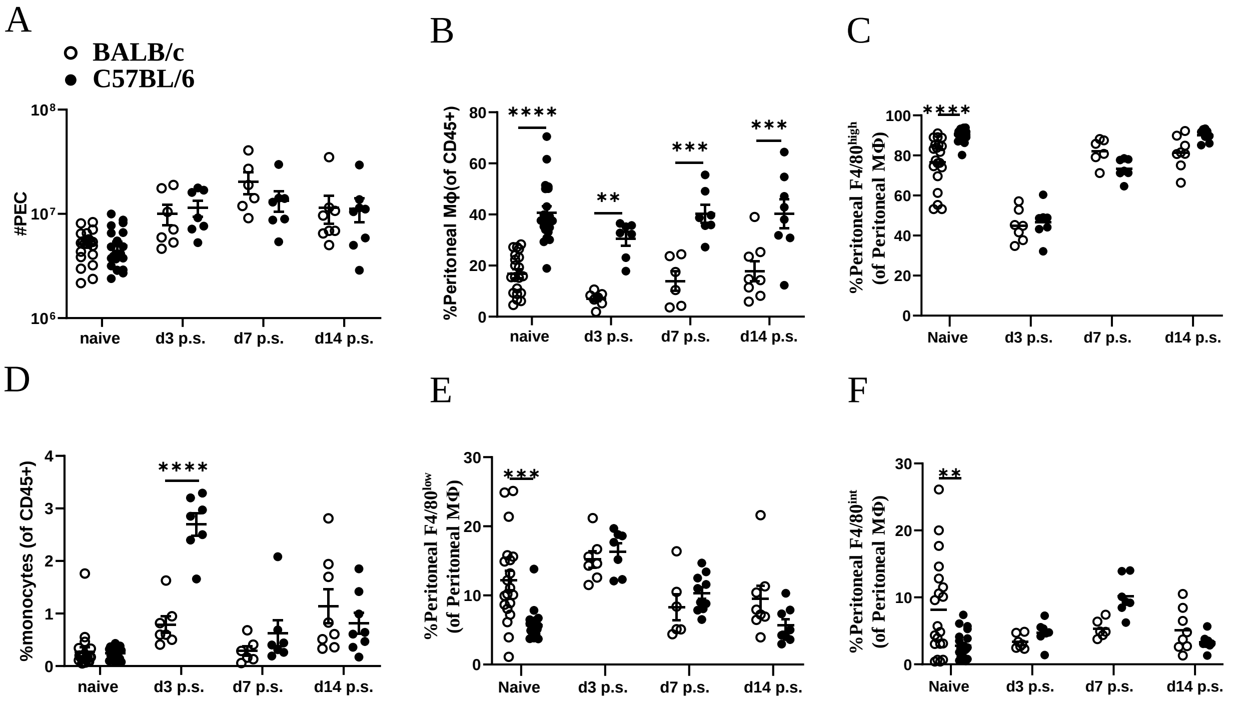


**Supplementary Figure 6.** Total Peritoneal exudate cells and relative percentage of peritoneal macrophages and monocytes in naïve mice or at various timepoints after surgery.

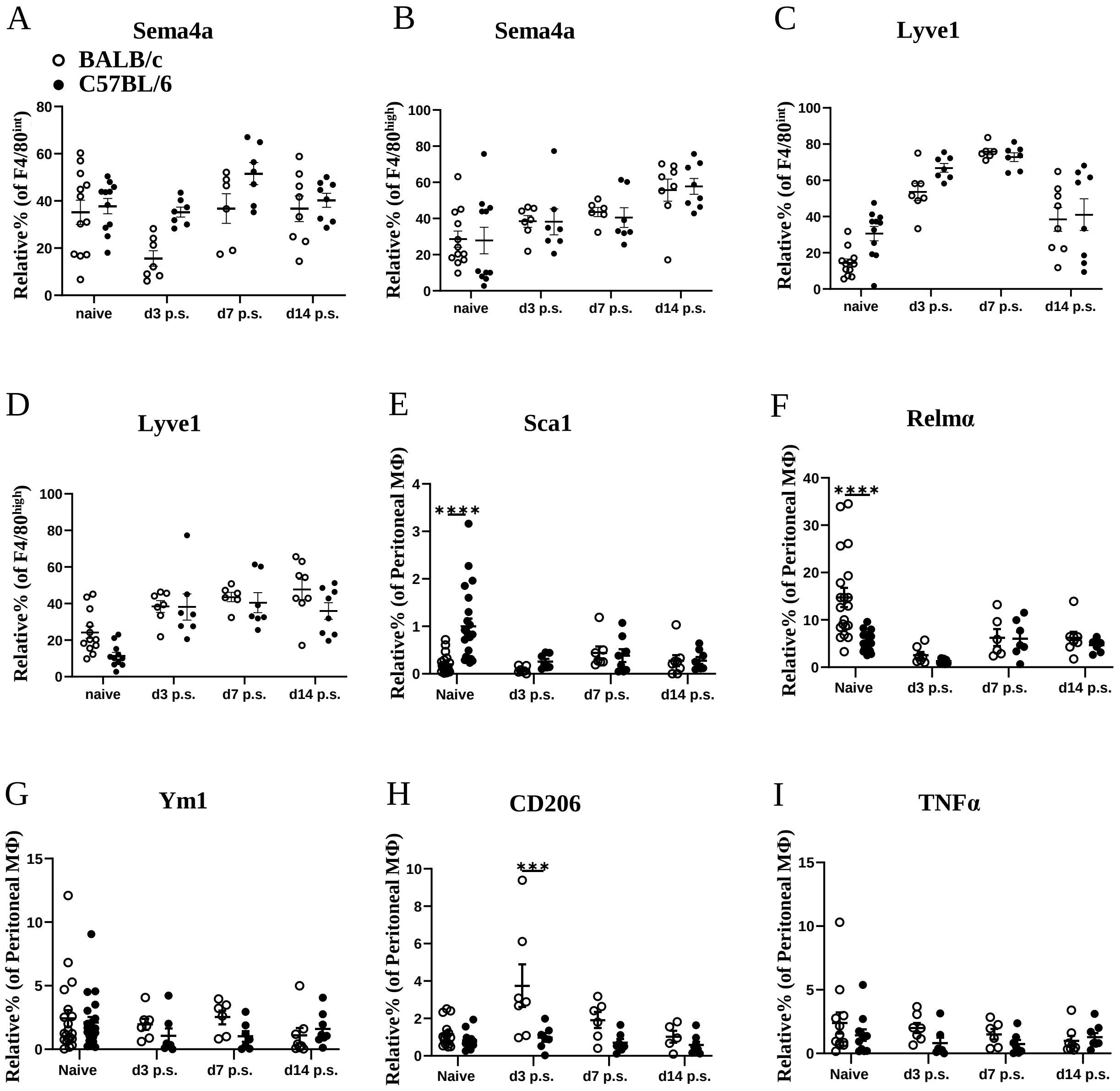


**Supplementary Figure 7.** Relative percentages for different markers expressed by peritoneal macrophages within BALB/c and C57BL/6 mice prior and post-surgery.

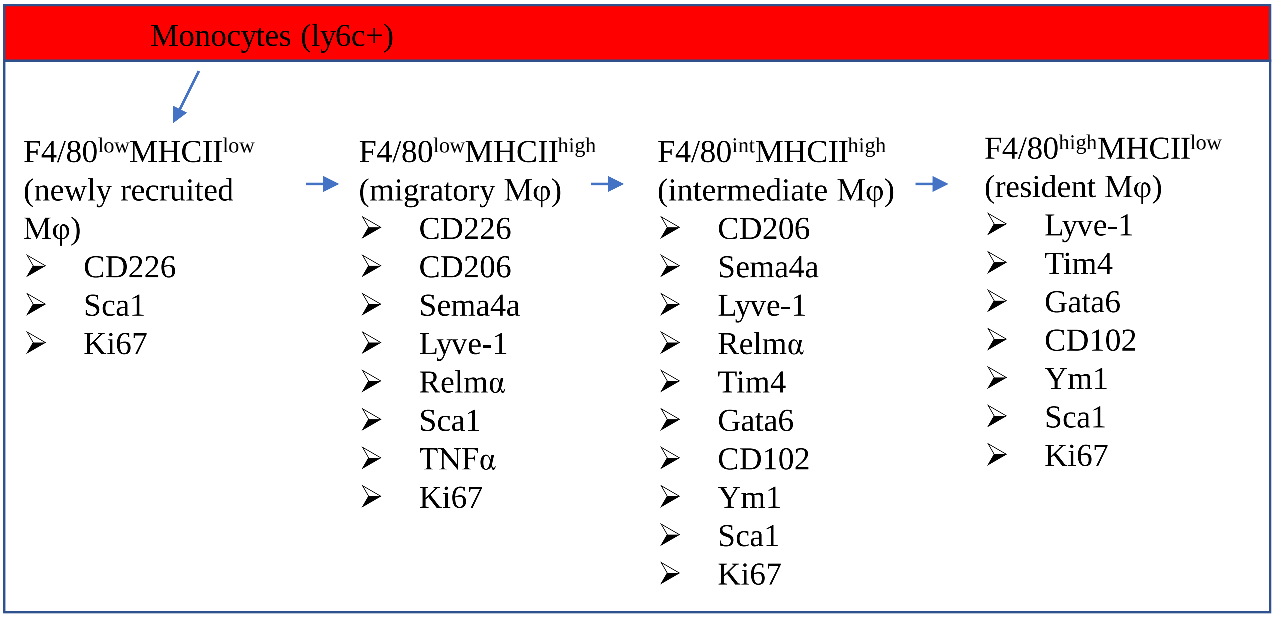


**Supplementary Figure 8.** Schematic representation of how different markers of peritoneal macrophages were expressed in different subsets

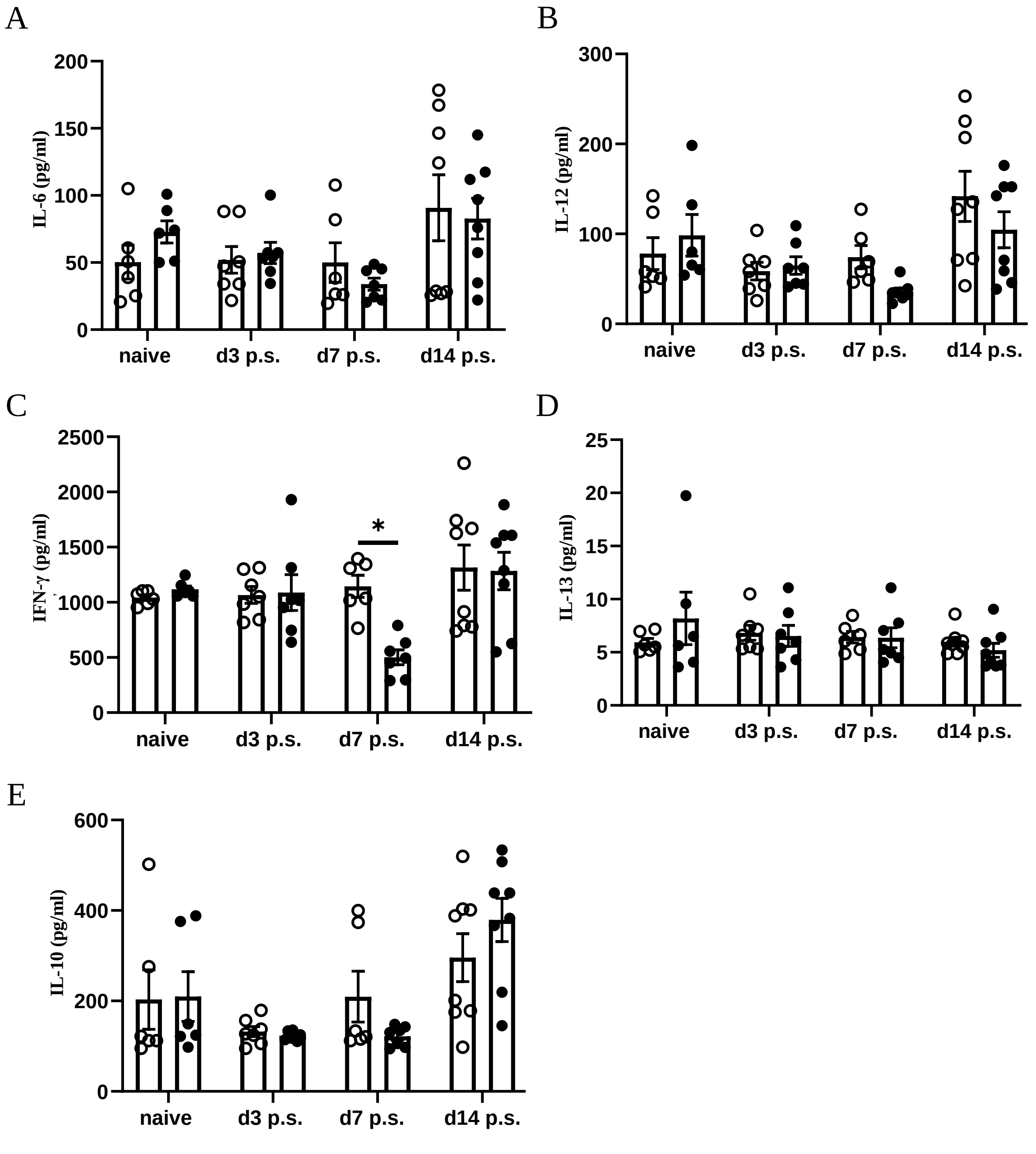


**Supplementary Figure 9.** Cytokine analysis in the peritoneal lavage of BALB/c and C57BL/6 before and after surgery.

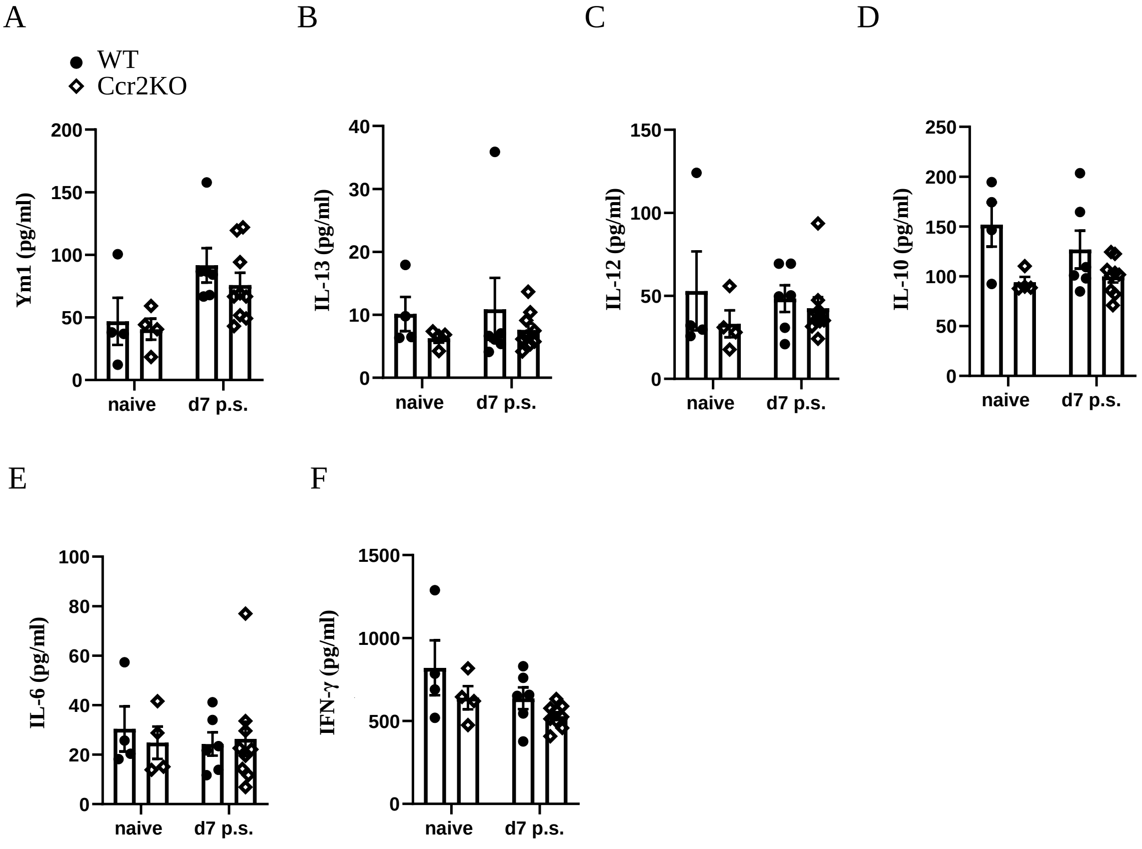


**Supplementary Figure 10.** Cytokine analysis in the peritoneal lavage of Ccr2KO and WT littermate before and after surgery.

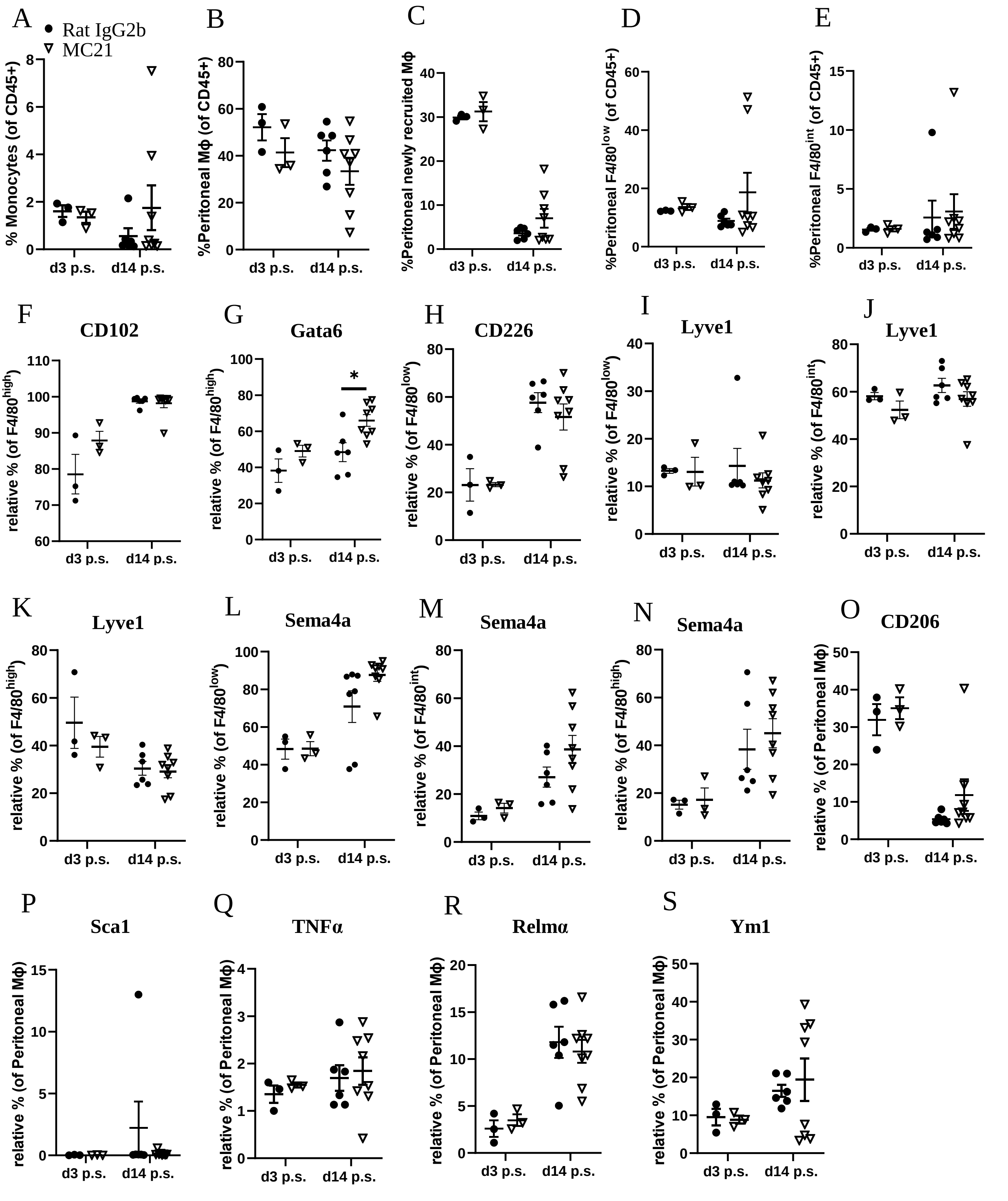


**Supplementary Figure 11.** Relative percentage of monocytes and different markers of peritoneal macrophages of Rat IgG2b isotype control and anti-CCR2 (clone MC21) treated mice after surgery. Peritoneal newly recruited macrophages are F4/80$^{low}$MHCII$^{low}$.

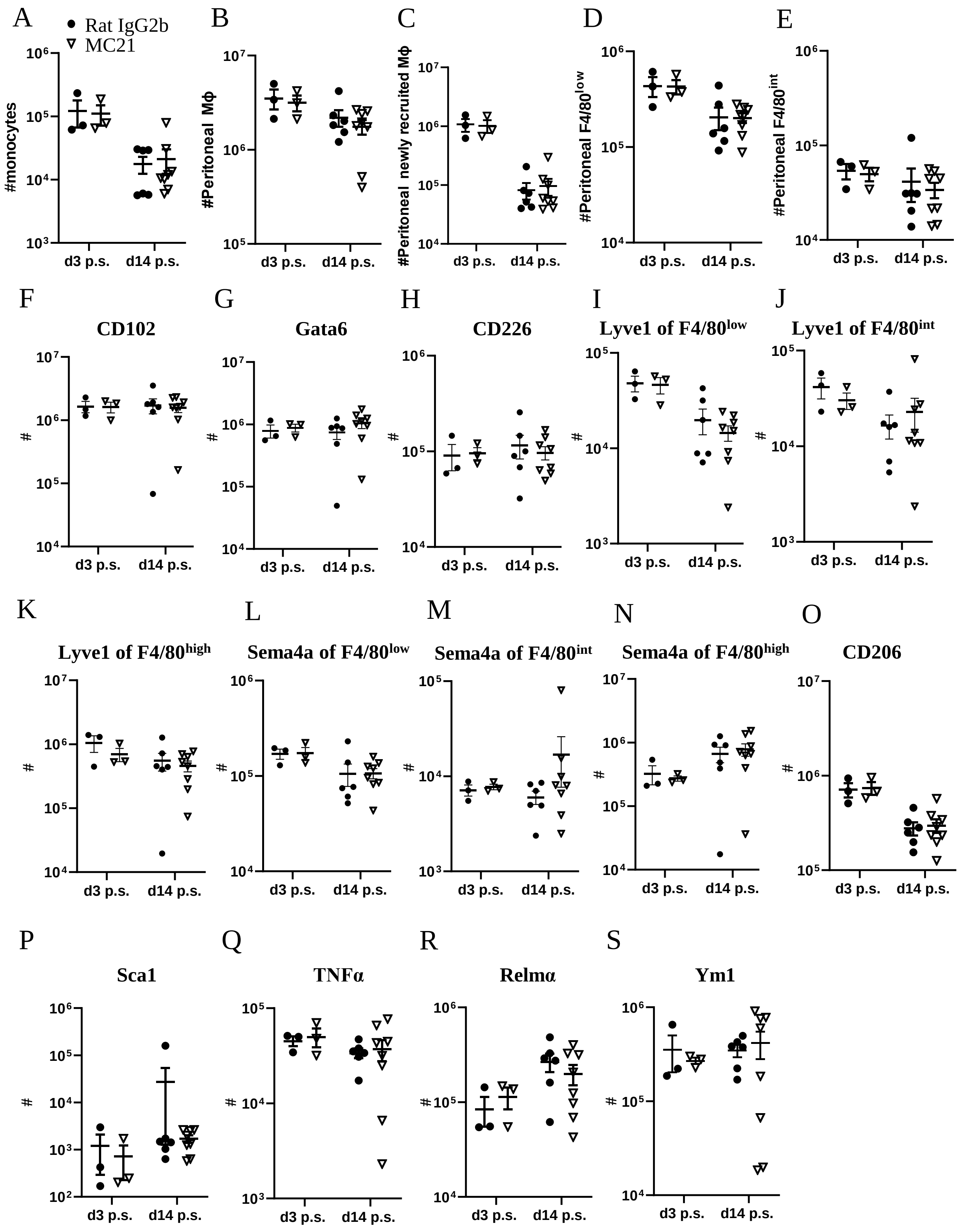


**Supplementary Figure 12.** Total cell number of monocytes and different markers of peritoneal macrophages of Rat IgG2b isotype control and anti-CCR2 (clone MC21) antibody treated mice after surgery. Peritoneal newly recruited macrophages are F4/80$^{low}$MHCII$^{low}$.

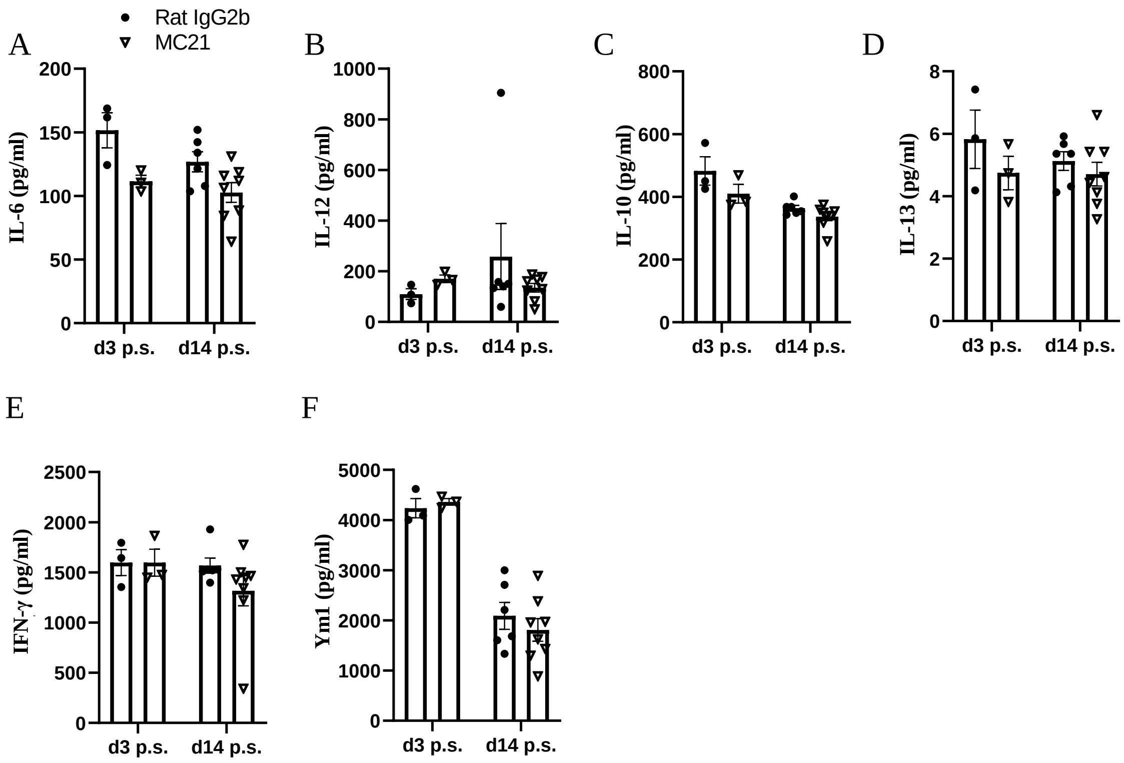


**Supplementary Figure 13.** Cytokine analysis in the peritoneal lavage of Rat IgG2b isotype control and anti-CCR2 (clone MC21) antibody treated mice after surgery.

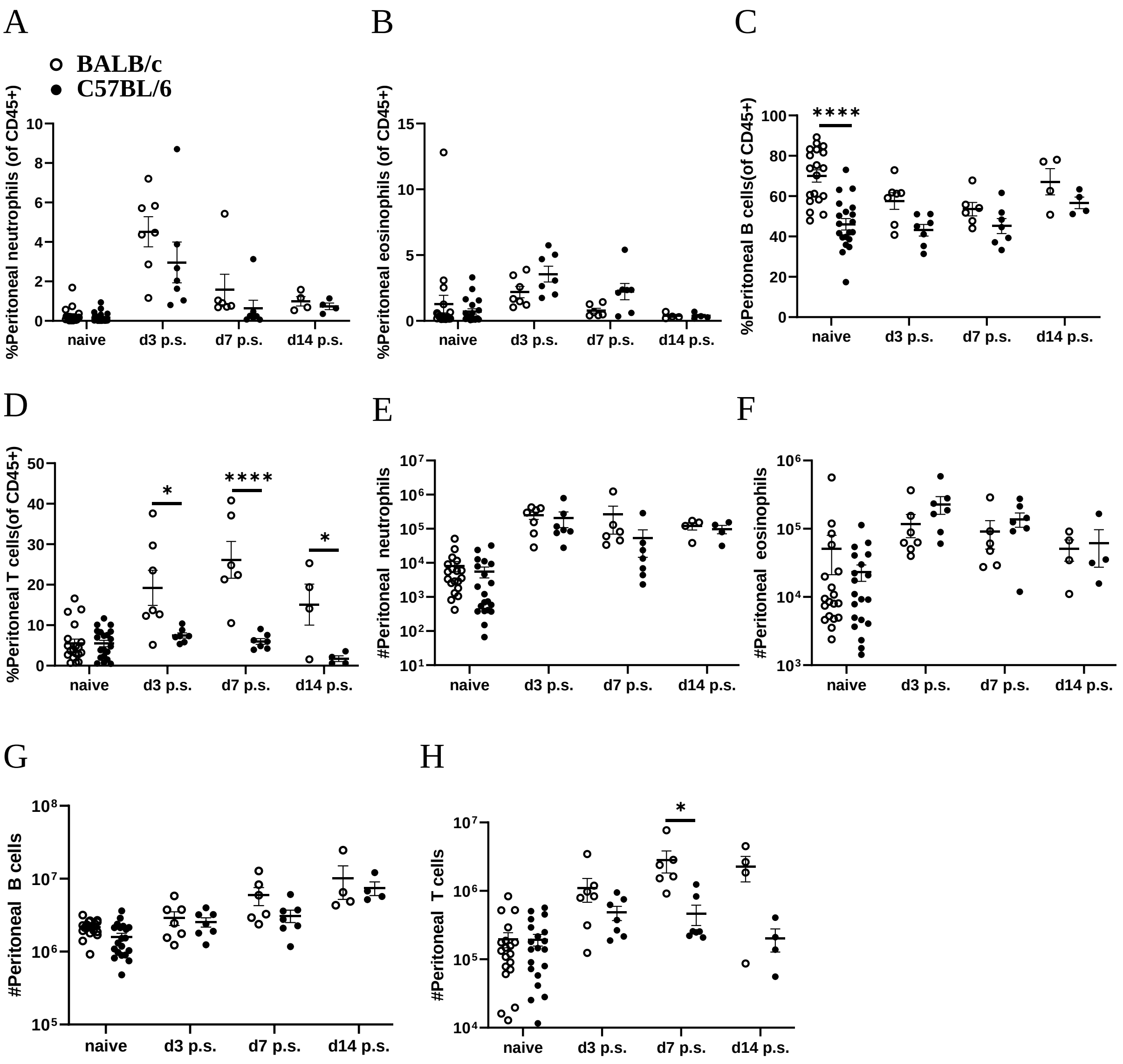


**Supplementary Figure 14.** The relative percentage and total cell number of neutrophils, eosinophils, B cells, and T cells in the peritoneal cavity of BALB/c and C57BL/6 mice prior and after surgery.
